# Neurocognitive signatures of intrinsic motivation in naturalistic information seeking and memory

**DOI:** 10.64898/2026.11.12.765917

**Authors:** Haowen Su, Coralie Knight, Philip R. Liu, Hongmi Lee

## Abstract

Humans often seek information without direct external reward. Theories distinguish two major dimensions of this intrinsic motivation: wanting, the desire to resolve an information gap, and liking, the enjoyment of the information itself. However, how these motivational dimensions shape self-guided information-seeking behavior, neural activity, and memory formation in real-world contexts remains underexplored. To address this gap, we developed a naturalistic fMRI paradigm in which participants freely browsed the internet, subsequently recalled their browsing experiences, and retrospectively reported their moment-to-moment wanting and liking. We found that both dimensions predicted more stable browsing trajectories but engaged partially dissociable neural systems: wanting recruited regions supporting goal-directed cognition, whereas liking engaged affective regions. Wanting and liking also interacted to predict memory, with wanting predicting recall primarily when liking was low. Compared with active browsing, passive viewing of the same web content reduced wanting and liking and diminished their dissociation, yet both continued to predict memory and intersubject neural similarity in the default mode network. These findings show that naturally fluctuating wanting and liking guide how we seek and remember information through overlapping yet distinct neurocognitive mechanisms.

## INTRODUCTION

In modern society, people spend much of each day seeking information, often by scrolling through webpages or browsing social media, even when these activities yield no tangible benefit. Consistent with this everyday behavior, empirical and theoretical work in psychology and neuroscience has long suggested that information itself can be intrinsically rewarding^1,2^. Humans and non-human animals seek information unrelated to external rewards and may even lead to punishment^3^, and such “curiosity-driven” information seeking engages the brain’s reward and valuation systems^3–7^. The intrinsic drive to seek information also supports learning and memory^2,8^, with numerous laboratory studies showing that curiosity at encoding enhances subsequent recall^4,5,9–13^. This benefit may arise through multiple mechanisms, including increased attention and frontoparietal activity^14,15^ and dopaminergic modulation of the medial temporal memory system^4,5,8,16^.

Although prior studies have typically conceptualized intrinsic motivation for information as a unitary anticipatory drive^4,5,17^, information seeking can arise from distinct motivational states^18–23^. Theoretical accounts highlight two largely independent dimensions^24,25^. One is “wanting”, or the desire to reduce uncertainty arising from a perceived information gap that is often accompanied by an aversive affective state^24,26,27^. The other is “liking”, or the enjoyment or interest elicited by the information itself, accompanied by a positive affective state^24,25,27^. While conceptually related, they are dissociable and may give rise to distinct forms of information seeking depending on their relative contributions^25^. For example, individuals may find information highly enjoyable without feeling compelled to seek more, as when savoring aesthetically pleasing content. Conversely, they may persist in seeking unpleasant information to resolve ambiguity, as in morbid curiosity^28^. These two dimensions can also differentially affect information-seeking behavior and memory performance^12,29–31^.

However, several questions remain about how wanting and liking influence information seeking and learning. First, the neural mechanisms underlying these two motivational dimensions are not fully understood. Although theoretical accounts associate wanting and liking with the dopaminergic and opioidergic systems, respectively^25,27^, direct comparisons of their neural correlates during information seeking are scarce. How the two dimensions dissociably or interactively shape neural responses is particularly underexplored, as previous neuroimaging studies have typically focused on a single construct, such as curiosity^4,5,17^. Second, the neurocognitive effects of wanting and liking have rarely been examined during self-guided information seeking in naturalistic settings^29,32,33^. Prior studies have predominantly used controlled paradigms in which participants passively engage with simple, experimenter-provided information, such as trivia questions^4,5,9,16^, with motivation assessed on discrete, randomized trials^12,30,31^. While these approaches have provided important evidence for the effects of intrinsic motivation, they may not fully capture how wanting and liking operate when individuals actively and continuously decide what information to pursue, or the broader range of motivational states that complex, self-relevant, real-world content may evoke.

To investigate how naturally fluctuating wanting and liking shape neurocognitive processes of information seeking and learning in real-world settings, we introduced a free web-browsing functional magnetic resonance imaging (fMRI) paradigm. Web browsing is a major form of information seeking and a primary source of knowledge in today’s world, and it is increasingly used to study information-seeking behavior^32–35^. Here, we extended this approach by combining self-guided browsing with fMRI and a subsequent memory test. Participants freely browsed the internet for 30 minutes and then verbally recalled their browsing experiences during scanning. They also retrospectively rated their moment-to-moment wanting and liking for the information they sought while watching a screen recording of their browsing session. This retrospective sampling captured changes in motivational states without interrupting the natural flow of browsing. Additionally, we included a passive-viewing condition in which independent participants watched recordings of others’ browsing sessions, allowing us to compare active information seeking with passive observation using comparable real-world content.

Using this design, we asked three questions: (1) How do wanting and liking relate to behavior and neural activity during naturalistic information seeking? (2) How do wanting and liking affect the behavioral and neural markers of successful memory formation? (3) How do active seekers and passive viewers differ in these two motivational dimensions and their relationship with memory? We found that wanting and liking were associated with more stable and semantically continuous information seeking, with liking particularly linked to reduced content-switching. The two dimensions showed overlapping but dissociable neural activation, with wanting more strongly associated with regions implicated in goal-directed cognition and liking with regions involved in affective processing. They also dissociated in their effects on memory: liking more consistently predicted recall and neural reinstatement, whereas wanting predicted memory primarily when liking was low. Finally, although passive viewers showed weaker and less differentiated wanting and liking than active seekers, their intrinsic motivation still predicted memory and neural responses shared across individuals. Together, these findings suggest that wanting and liking shape naturalistic information seeking and memory through partially shared but functionally distinct mechanisms, providing a more nuanced understanding of how intrinsic motivation guides learning in ecologically valid, real-world contexts.

## RESULTS

### Intrinsic motivation and behavioral patterns during active web browsing

We first examined how naturally fluctuating intrinsic motivation shapes information-seeking behavior during active web browsing. During fMRI, participants freely browsed the internet for 30 minutes using an MR-compatible mouse and then recalled what they remembered from their browsing experience (Figs. 1a-b). Participants also retrospectively rated their moment-to-moment wanting and liking for information encountered during their browsing, while watching a screen recording of their activity (Fig. 1c). To characterize continuous browsing behavior, independent annotators segmented each browsing session into discrete events, each representing a coherent unit of content (e.g., an article or social media post), which served as the primary unit of analysis. Event-level measures of intrinsic motivation and browsing actions were derived by averaging wanting and liking ratings, on-screen keypress, click, and scroll frequencies, and mouse cursor distance traveled across all time points within each event (Fig. 2b). Across participants, 283 unique websites were visited (Fig. 2a). On average, each participant visited 10.11 websites (SD = 3.59) per session, comprising 140.6 events (SD = 36.6), with each event lasting 13.91 s (SD = 3.43). Supplementary Table 1 presents descriptive statistics for the individual motivation and browsing action measures.

**Fig. 1.**
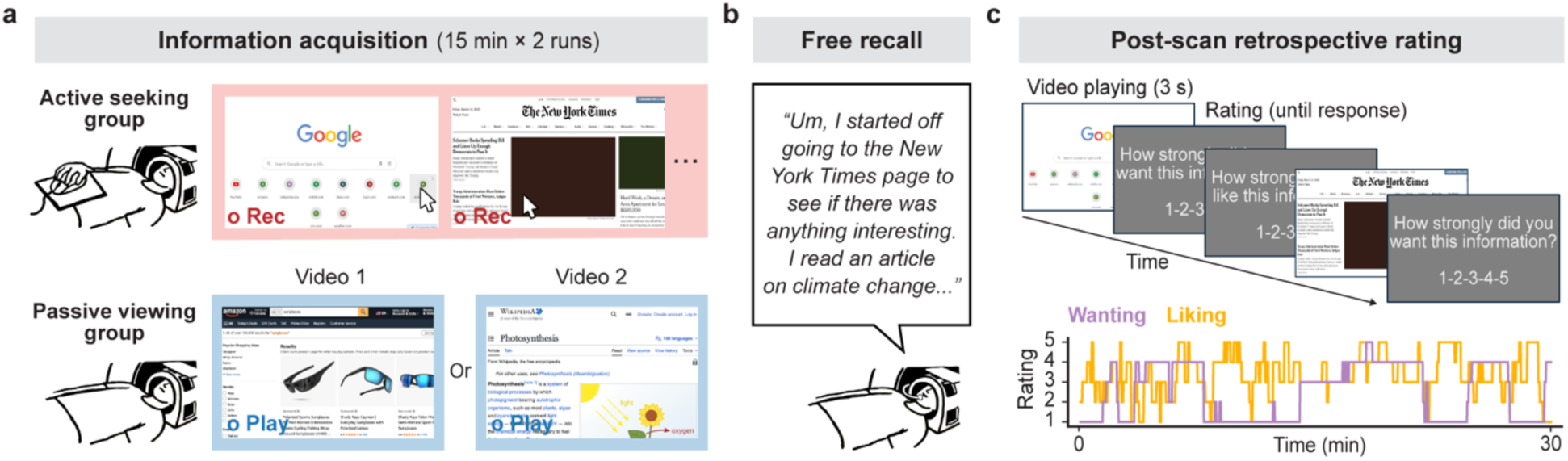
Study procedures. (**a**) Information acquisition phase. Participants were assigned to complete two 15-minute fMRI scanning runs of either the active information-seeking task or the passive viewing task. Participants in the active-seeking group freely browsed online information while their screen activity was recorded (top panel). Participants in the passive-viewing group watched screen-recording videos of web browsing performed by one of two selected active seekers (bottom panel). Red “Rec” labels indicate screen recordings from the active-seeking condition, whereas blue “Play” labels indicate video playback during the passive-viewing condition. These labels were added for illustration purposes and were not visible to participants during the actual experiment. (**b**) Free recall phase. Participants verbally recalled their experiences during the information acquisition phase while their responses were audio-recorded. (**c**) Post-scan retrospective rating phase. Participants rated their moment-to-moment wanting and liking during the information acquisition phase on a 1– 5 Likert scale while watching a screen recording of the web-browsing session they had performed or observed (top panel). The resulting ratings produced continuous time series for each of the two motivational dimensions, illustrated by one example participant’s ratings (bottom panel).

**Fig. 2.**
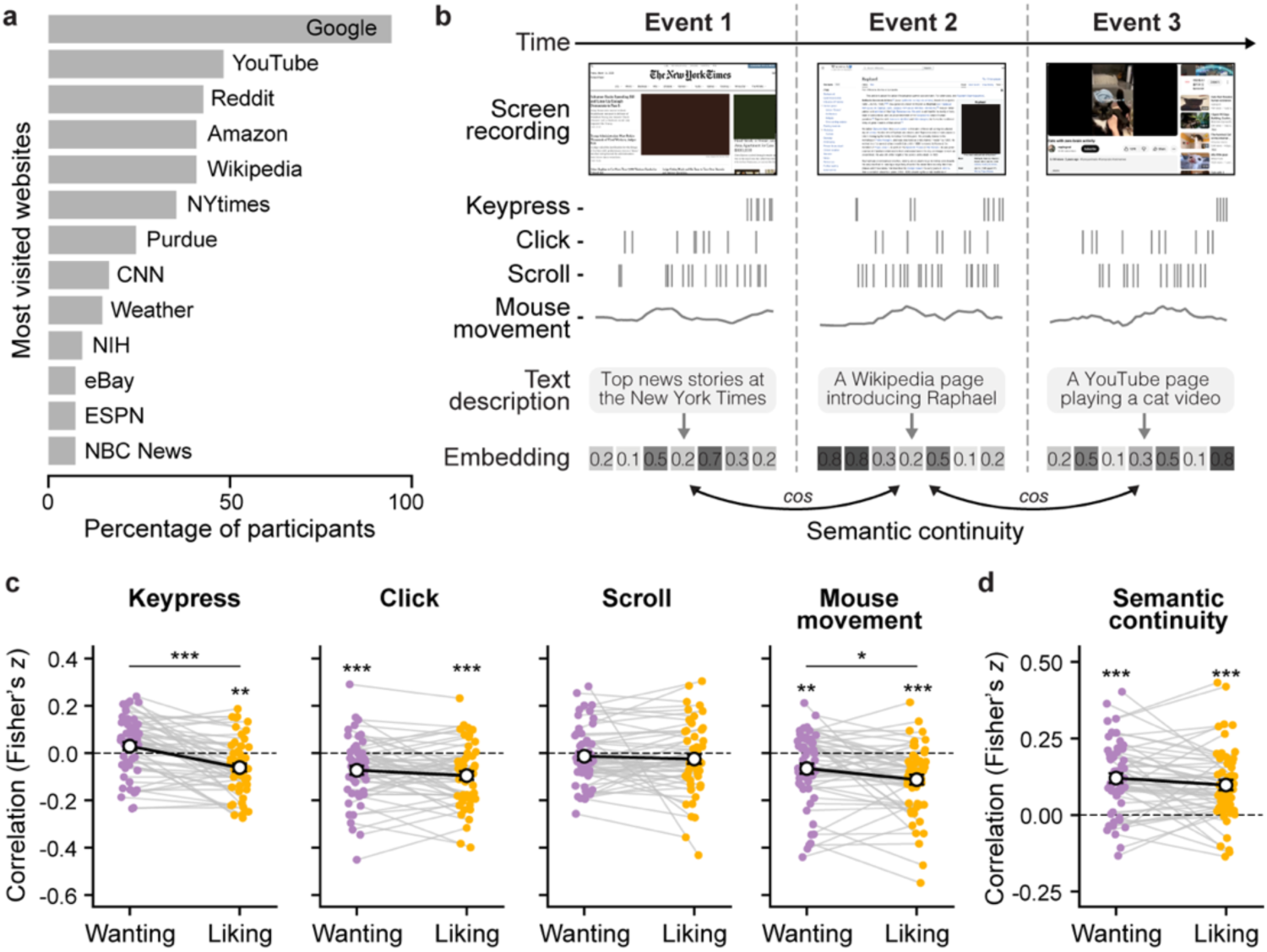
Behavioral characteristics of active information seeking through web browsing. (**a**) Most frequently visited websites. Bar graphs show the percentage of participants in the active information-seeking group who visited each website (domain) at least once during the web browsing session. (**b**) Event-level feature extraction. Web browsing screen recordings were manually segmented into events based on changes in content. Browsing actions were averaged within each event, including click, scroll, and keypress frequencies, as well as the distance traveled by the mouse cursor (mouse movement). Human annotators generated text descriptions of the content of each event, which were then embedded using a language model to compute semantic continuity (cosine similarities) between adjacent events. (**c**) Correlations between mean intrinsic motivation (wanting and liking) and browsing actions across events within each participant. (**d**) Correlations between mean intrinsic motivation and semantic continuity to the subsequent event. In **c** and **d**, colored dots represent individual participants, and white circles indicate group means. Significance relative to zero was assessed using two-tailed one-sample *t*-tests for wanting and liking separately. Differences between wanting and liking were assessed using two-tailed paired *t*-tests. Error bars indicate the SEM across participants. \**p* < 0.05, \*\**p* < 0.01, \*\*\**p* < 0.001 (uncorrected).

To examine how intrinsic motivation relates to information-seeking behavior, we tested associations between each type of browsing action and ratings of wanting and liking across events. Within-participant Pearson correlations revealed that both wanting and liking were negatively associated with click frequency and mouse movement, while liking was additionally negatively associated with keypress frequency (all *ps* < 0.01; Fig. 2c, Supplementary Table 2). Direct comparisons between wanting and liking further revealed that liking showed a stronger negative association with keypress frequency (*t*(51) = 5.40, *p* < 0.001, 95% CI = [0.06, 0.12], Cohen’s *d* = 0.73) and mouse movement (*t*(52) = 2.44, *p* = 0.018, 95% CI = [0.01, 0.08], Cohen’s *d* = 0.30) than wanting did. Thus, higher wanting and liking were generally accompanied by fewer actions associated with content-switching and searching, with this pattern more pronounced for liking than for wanting.

We then asked whether intrinsic motivation shaped the trajectory of naturalistic information seeking, specifically whether participants tended to transition to semantically related content. To quantify semantic continuity between successive browsing events, human annotators described each event in text based on the screen recordings. Semantic continuity was then computed as the cosine similarity between the sentence embeddings of the two events’ descriptions (Fig. 2b). Event-wise motivation ratings correlated positively with semantic continuity to the following event for both wanting (*t*(52) = 7.45, *p* < 0.001, 95% CI = [0.09, 0.15], Cohen’s *d* = 1.02) and liking (*t*(53) = 6.09, *p* < 0.001, 95% CI = [0.07, 0.13], Cohen’s *d* = 0.83), with no significant difference between the two dimensions (*t*(52) = 1.46, *p* = 0.149, 95% CI = [−0.01, 0.05], Cohen’s *d* = 0.19; Fig. 2d). This suggests that higher intrinsic motivation was followed by a shift to semantically related content.

### Brain activity associated with intrinsic motivation during active web browsing

We next examined the neural correlates of intrinsic motivation during active web browsing. First, we performed a whole-brain linear mixed-effects regression analysis predicting event-level activation in each cortical parcel of the Schaefer 200-parcel atlas^36^ from motivation ratings. Because wanting and liking were positively correlated within participants (mean *r* = 0.51, SD = 0.25), we initially modeled each motivational dimension separately, with participant as a random effect. All models included event-level browsing actions, low-level perceptual features of the screen recording, and scanning run as covariates.

The group-level activation maps revealed that wanting and liking were associated with distributed, partially overlapping sets of brain regions (Fig. 3a, Supplementary Fig. 2, Supplementary Tables 3-4). Both dimensions were related to greater activation in lateral and ventral temporal regions involved in high-level visual and semantic processing^37,38^ and to reduced activation in somatomotor regions. Despite this overlap, the two dimensions also showed dissociable patterns. Greater lateral prefrontal activation, for example, was significantly associated with higher wanting but not liking, whereas greater activation in the insula and anterior cingulate cortex, regions implicated in negative affect and cognitive conflict^39,40^, was significantly associated with lower liking but not wanting. Analyses of a priori subcortical regions implicated in motivation and memory formation^8,41^ (hippocampus, amygdala, caudate, and putamen; Supplementary Fig. 3) likewise showed overlapping but dissociable associations with wanting and liking (Supplementary Table 5). Hippocampal and amygdala activation were positively related to both wanting and liking (all *p*s < 0.05), whereas caudate activation was positively related to wanting only (*β* = 0.006, *t* = 2.18, *p* = 0.029, 95% CI = [0.001, 0.010]).

**Fig. 3.**
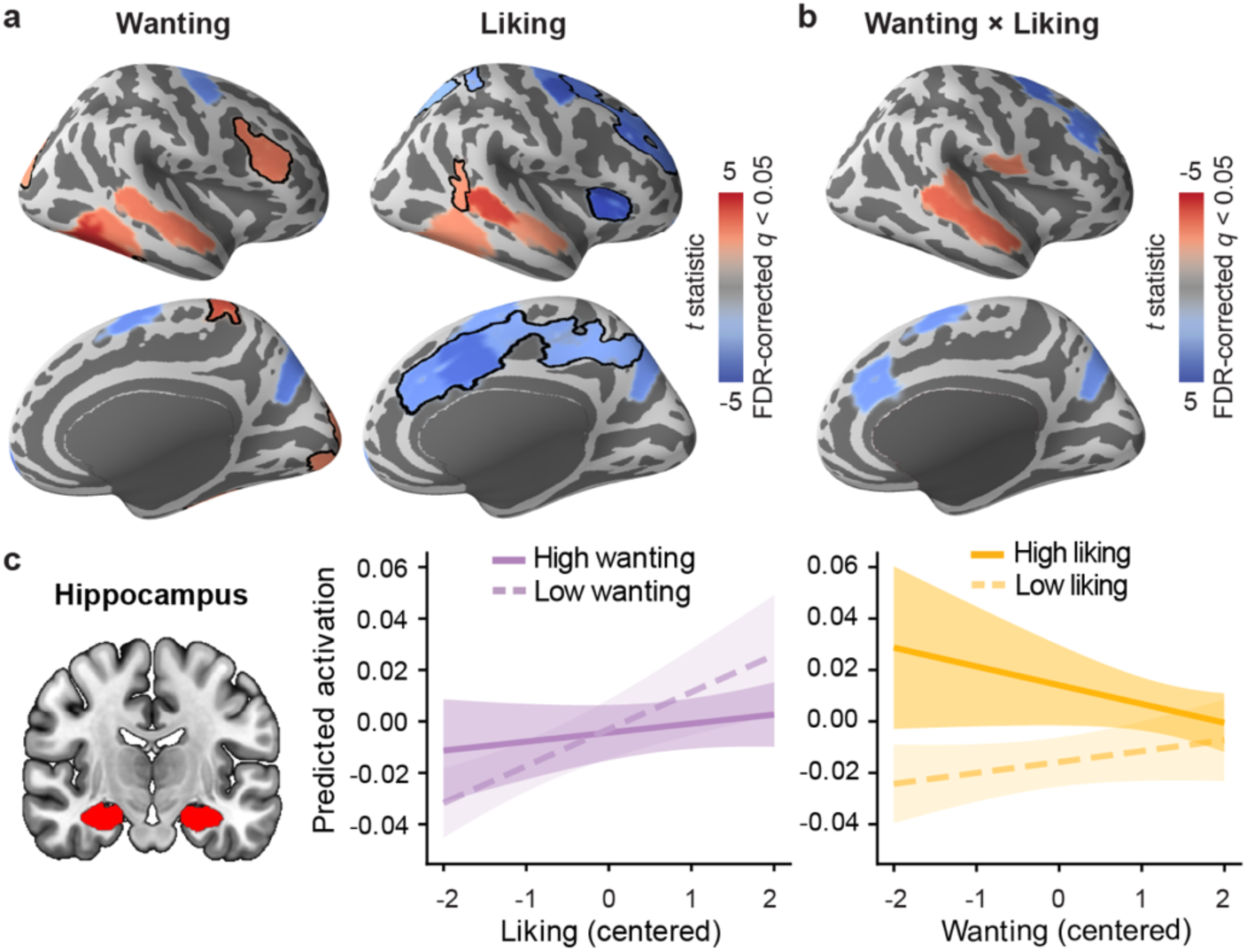
Neural activation associated with intrinsic motivation during web browsing. (**a**) Cortical regions whose activation was associated with wanting (left) and liking (right). Red and blue regions indicate positive and negative associations with the motivational dimension of interest, respectively. Black outlines denote regions that were uniquely significant for the corresponding motivational dimension. Statistical significance was assessed for each cortical parcel using separate linear mixed-effects models relating event-wise mean neural activation to either wanting or liking. (**b**) Cortical regions whose activation was associated with the interaction between wanting and liking, after controlling for their main effects. Red regions indicate relatively more positive effects of wanting when liking is lower. Blue regions indicate relatively more positive effects of wanting when liking is higher. In **a** and **b**, *t*-statistic maps are visualized on the inflated lateral (top) and medial (bottom) surfaces of the right hemisphere of the FreeSurfer template brain (fsaverage). Maps were thresholded at *q* < 0.05 (FDR-corrected across all cortical parcels). Full statistics for individual parcels are provided in Supplementary Tables 3, 4, and 6. (**c**) Interactive effects of wanting and liking on bilateral hippocampal activation. The interaction between wanting and liking was tested using a mixed-effects model. Simple slopes were estimated at low and high levels (±1 SD from the mean) of wanting (middle panel) and liking (right panel), shown as dotted and solid lines, respectively. Shaded areas indicate 95% confidence intervals. Full statistics for the hippocampus and additional subcortical regions are provided in Supplementary Table 7.

We then fit models with wanting, liking, and their interaction as simultaneous predictors to estimate the unique association of each dimension with neural activity (Supplementary Fig. 4; Supplementary Tables 3, 6-7). Consistent with the separate models, wanting remained positively associated with activation in the lateral prefrontal cortex and caudate (all *p*s < 0.04), whereas liking remained negatively associated with activation in the insula and anterior cingulate cortex and positively associated with activation in the hippocampus and amygdala (all *p*s < 0.01). Wanting and liking also interacted in several regions, including some associated with both dimensions in the separate models (Figs. 3b-c), such as the superior temporal sulcus (right TempPar_1; *β* = −0.01, *t* = −3.65, *p* < 0.001, 95% CI = [−0.012, −0.004]) and hippocampus (*β* = −0.005, *t* = −2.49, *p* = 0.013, 95% CI = [−0.008, −0.001]). Simple slope analyses showed that the association between wanting and activation in these regions was more positive at low than high levels of liking (Supplementary Table 8). Overall, wanting and liking showed partially distinct neural associations, with wanting more strongly related to regions implicated in goal-directed cognition and liking to regions implicated in affective processing. Their interaction also suggests that the neural correlates of one dimension may depend on the level of the other, consistent with accounts proposing that different combinations of wanting and liking give rise to distinct motivational states during information seeking^25^.

### Recall performance for active web browsing experiences

Having shown that wanting and liking differentially relate to information-seeking behavior and neural activity during web browsing, we next asked whether they also relate to participants’ narrated recall of their browsing. On average, recall speech lasted 10.82 min (SD = 6.64) and contained 1395.2 words (SD = 939.73 words). To quantify recall performance, independent annotators coded each browsing event as remembered or forgotten by matching participants’ transcribed recall to the screen recordings (Fig. 4a). Participants recalled 59% of events on average (SD = 0.16). Mean motivation ratings (Fig. 4b) were higher for subsequently remembered than forgotten events for both wanting (*t*(53) = 2.82, *p* = 0.007, 95% CI = [0.05, 0.32], Cohen’s *d* = 0.38) and liking (*t*(53) = 4.63, *p* < 0.001, 95% CI = [0.16, 0.41], Cohen’s *d* = 0.63), consistent with prior evidence linking intrinsic motivation to enhanced memory^4,8,31^.

**Fig. 4.**
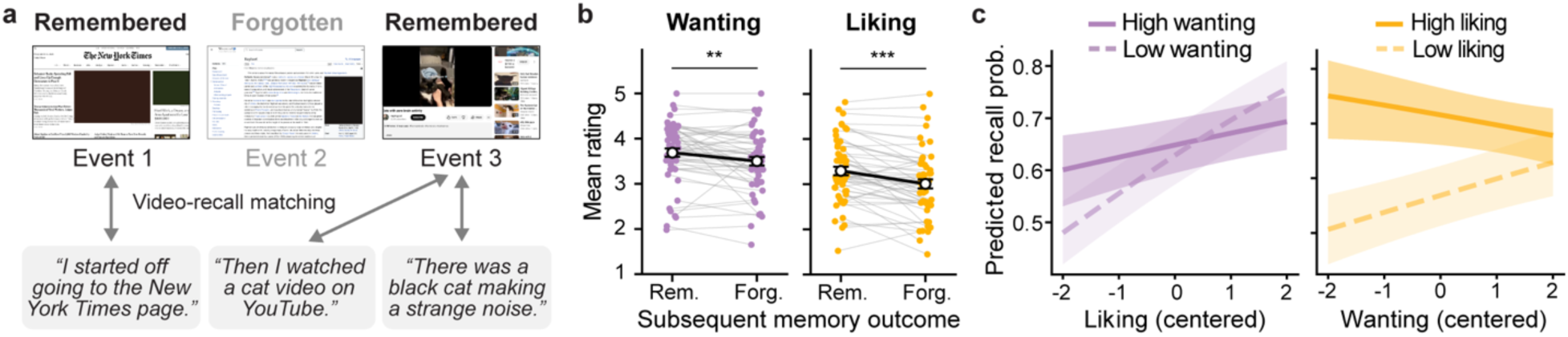
Intrinsic motivation and memory for web browsing experiences. (**a**) Alignment of free recall narratives with web browsing events. Each sentence in a participant’s recall transcript was manually matched to the corresponding events in that participant’s screen recording video. Events that were not mentioned during recall were classified as forgotten. (**b**) Mean wanting and liking ratings for subsequently remembered (Rem.) and forgotten (Forg.) events. Colored dots represent individual participants, and white circles indicate group means. Error bars indicate the SEM across participants. Two-tailed paired *t*-tests revealed that both wanting and liking ratings were higher for remembered than for forgotten events. \*\**p* < 0.01, \*\*\**p* < 0.001. (**c**) Interactive effects of wanting and liking on the probability of subsequent recall. The interaction was tested using a logistic mixed-effects model. Simple slopes were estimated at low and high levels (±1 SD from the mean) of wanting (left panel) and liking (right panel), shown as dotted and solid lines, respectively. Shaded areas indicate 95% confidence intervals.

We further tested whether wanting, liking, and their interaction made unique contributions to event-level memory, using a mixed-effects logistic regression that controlled for browsing actions and low-level perceptual features (Fig. 4c). Liking positively predicted subsequent recall (*β* = 0.26, *z* = 7.79, *p* < 0.001, 95% CI = [0.19, 0.32]), whereas the main effect of wanting was not significant (*β* = 0.04, *z* = 1.17, *p* = 0.241, 95% CI = [−0.02, 0.10]). The wanting-by-liking interaction was also significant (*β* = −0.09, *z* = −4.41, *p* < 0.001, 95% CI = [−0.12, −0.05]). Simple slopes analyses (Supplementary Table 9) showed that liking positively predicted recall at both low and high levels of wanting, whereas wanting did so only at low levels of liking (all *ps* < 0.005). Together with the wanting-by-liking interactions in neural activation reported above, these results suggest that different combinations of wanting and liking may reflect distinct motivational states with different mnemonic consequences. For example, unexpectedly pleasant content may be particularly memorable: events with low wanting (mean rating < 2.5) and high liking (mean rating > 3.5) had the numerically highest recall probability (M = 0.65, SD = 0.38), followed by high wanting and high liking (M = 0.63, SD = 0.22), high wanting and low liking (M = 0.58, SD = 0.32), and low wanting and low liking (M = 0.50, SD = 0.31).

### Neural markers of memory for active web browsing experiences

Wanting and liking during web browsing were related to behavioral memory performance. Were they also linked to neural markers of memory encoding and recall? We first identified brain regions whose activation during web browsing predicted subsequent recall, using a whole-brain mixed-effects logistic regression. Group-level results (Fig. 5a, Supplementary Table 10) showed activation in regions that largely overlapped with those related to intrinsic motivation, particularly liking. Specifically, successful recall was associated with lower activation in the insula, anterior cingulate cortex, and somatomotor regions, but higher activation in the lateral temporal cortex. Amygdala activation also predicted recall success, and hippocampal activation showed a similar but nonsignificant trend (Supplementary Table 11).

**Fig. 5.**
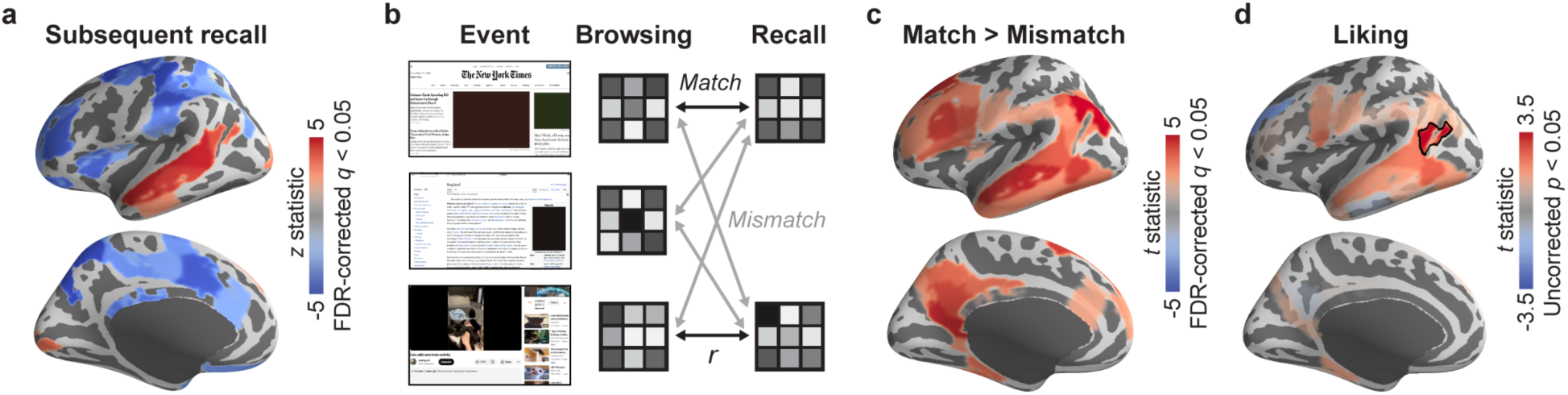
Neural markers of web browsing memory formation and recall. (**a**) Cortical regions whose activation during a web browsing event was associated with subsequent recall of the event. Red and blue regions indicate higher activation predicting subsequent remembering and forgetting, respectively. (**b**) Encoding-retrieval multivoxel pattern similarity analysis. For each cortical parcel, its event-wise multivoxel activation patterns (grayscale grids, with each square representing a voxel and shades representing activation levels) were correlated across the web browsing and recall phases. The browsing-recall event pairs were either matching (same events) or non-matching (different events). (**c**) Cortical regions showing event-specific memory reinstatement during recall. Red regions indicate higher browsing-recall pattern similarity for matching than non-matching web browsing event pairs. In **a** and **c**, z– and *t*-statistic maps are thresholded at *q* < 0.05 (FDR-corrected across all cortical parcels). Full statistics for individual parcels are provided in Supplementary Tables 10 and 12. (**d**) Cortical regions whose encoding-retrieval pattern similarity was associated with liking. Red and blue regions indicate higher liking during web browsing predicting higher and lower encoding-retrieval pattern similarity, respectively. Statistical maps of *t* values are thresholded at *p* < 0.05 (uncorrected). Black outlines denote parcels that survived FDR correction among all parcels included in the analysis. In **a**, **c**, and **d**, z– and *t*-statistic maps are visualized on the inflated lateral (top) and medial (bottom) surfaces of the left hemisphere.

Next, we performed an encoding-retrieval pattern similarity analysis^42^ to test whether intrinsic motivation was associated with neural reinstatement of browsing experiences during recall. For each cortical parcel, reinstatement was quantified as the correlation between multivoxel activation patterns during browsing and recall for corresponding events (Fig. 5b). Consistent with prior findings^43,44^, we observed event-specific reinstatement, reflected in greater pattern similarity between matching than non-matching events, in default mode network regions^45,46^ including posterior medial, lateral parietal, and lateral temporal cortices (Fig. 5c, Supplementary Table 12). In these regions, linear mixed-effects models with wanting, liking, and their interaction as predictors showed that liking was positively related to neural reinstatement in the left superior temporal gyrus (Fig. 5d; *β* = 0.014, *t* = 3.45, *p* < 0.001, 95% CI = [0.01, 0.02]). Neither wanting nor the interaction was significantly associated with reinstatement in any region (all *t*s < 1.85, *q*_FDR_ > 0.05). These results demonstrate robust reinstatement of browsing experiences during recall, and suggest that this reinstatement tracks liking more reliably than wanting.

### Motivation and memory during active web browsing versus passive viewing

Thus far, we have examined how intrinsic motivation relates to web browsing and subsequent memory in the active information-seeking condition, in which participants fully controlled what information to access. Information can also be acquired through passive exposure, however, and prior studies have largely examined passive acquisition using experimenter-selected content^4,5,47^. We therefore tested whether active browsing differed from passive viewing in intrinsic motivation during information acquisition and subsequent memory. In the passive-viewing condition, independently recruited participants (viewers) were randomly assigned to watch one of two screen recordings (Video 1 or Video 2) generated by participants in the active information-seeking condition (seekers). They then completed the same recall task and motivation ratings (Fig. 1). The seeker and viewer groups, as well as the two viewer groups assigned to different videos, did not differ significantly in demographic or personality measures (Supplementary Tables 13-14).

We first compared intrinsic motivation between the active information-seeking and passive-viewing conditions (Figs. 6a-b, Supplementary Tables 15-16). At the group level, Welch’s *t*-tests showed that seekers reported significantly higher average wanting than both Video 1 and Video 2 viewers, and significantly higher average liking than Video 2 viewers (all *p*s < 0.001). Seekers also showed a weaker within-participant correlation between wanting and liking than both viewer groups (*p*s < 0.01), indicating greater independence between the dimensions. One-sample *t*-tests comparing viewers with the seeker who generated the video they watched revealed the same pattern, except that the wanting-liking correlation did not differ significantly between Video 2 viewers and their seeker (*p* = 0.161). Thus, active seeking was associated with higher intrinsic motivation, particularly wanting, and potentially greater diversity in motivational states, characterized by different combinations of wanting and liking.

**Fig. 6.**
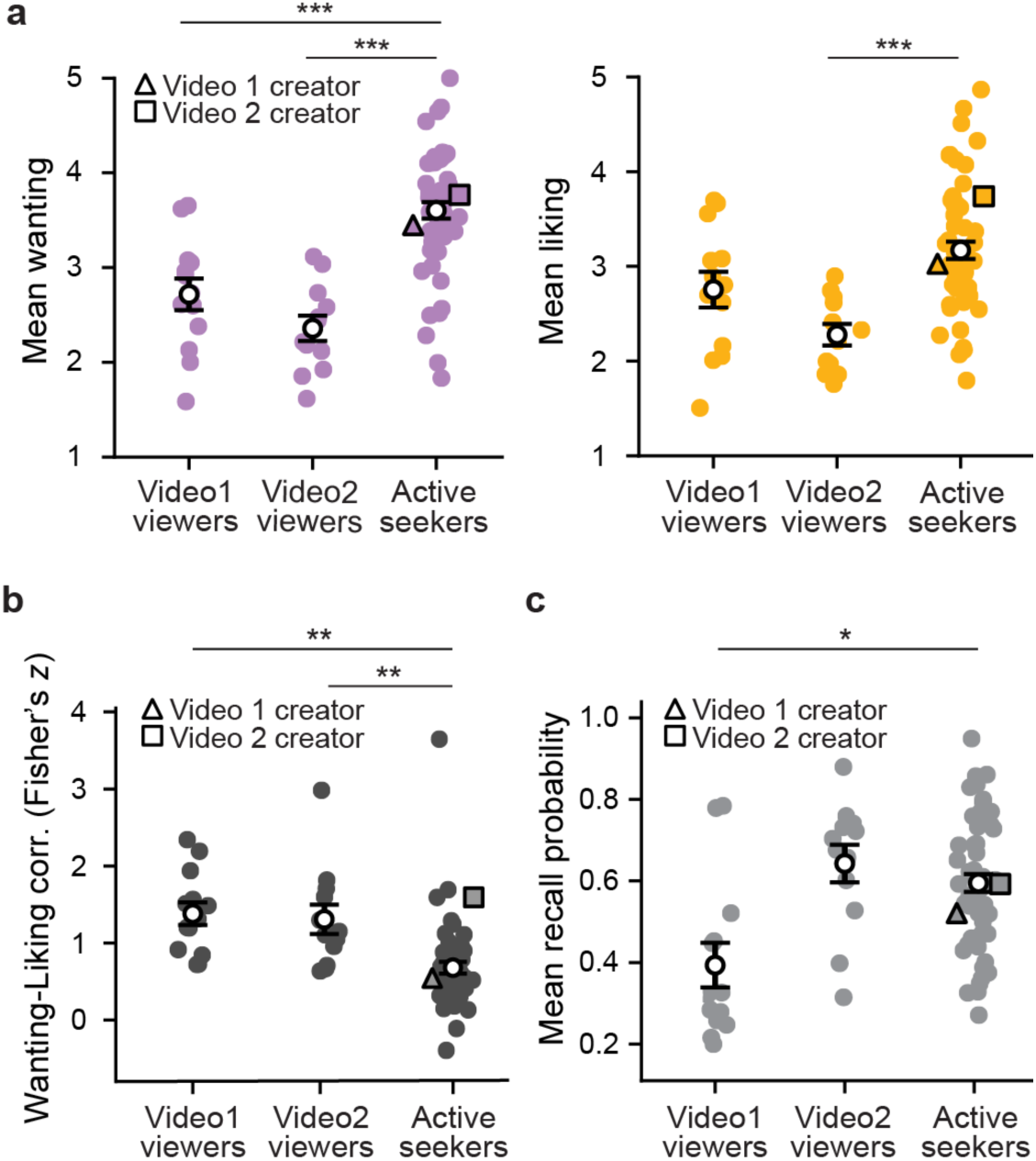
Comparison of intrinsic motivation and memory between passive viewers and active information seekers. (**a**) Mean wanting (left panel) and liking (right panel) ratings for viewers while watching Video 1 or Video 2 and for seekers during active web browsing. (**b**) Within-participant correlations between event-wise wanting and liking ratings in viewers and seekers. (**c**) Mean recall probability of events in viewers and seekers. In **a**-**c**, colored symbols represent individual participants, and white circles indicate group means. Triangles (Video 1) and squares (Video 2) denote the two seekers whose screen recording videos were used as stimuli in the passive-viewing condition. Error bars indicate the SEM across participants. Group differences were assessed using two-tailed Welch’s independent-samples *t*-tests. \**p* < 0.05, \*\**p* < 0.01, \*\*\**p* < 0.001.

We then compared memory performance between the active-seeking and passive-viewing conditions (Fig. 6c, Supplementary Tables 15-16). The mean proportion of recalled events was higher among seekers than among Video 1 viewers (*t*(12.47) = 2.86, *p* = 0.014, 95% CI = [0.05, 0.34], Hedges’ *g* = 0.96), but not Video 2 viewers (*p* = 0.340). Recall also did not differ significantly between viewers and the seeker whose recording they watched, in either video group (*p*s > 0.09). Thus, although seekers showed higher overall intrinsic motivation, this did not consistently translate into better recall. To test whether this was because motivation was unrelated to memory during passive viewing, we fit mixed-effects logistic regressions predicting viewers’ event-level recall from motivation ratings. Because wanting and liking were highly correlated during passive viewing, separate models were fit for each dimension. Both wanting (*β* = 0.24, *z* = 6.78, *p* < 0.001, 95% CI = [0.17, 0.31]) and liking (*β* = 0.15, *z* = 4.23, *p* < 0.001, 95% CI = [0.08, 0.23]) positively predicted recall, indicating that the motivation-memory association was present during passive viewing as well as active seeking.

### Shared memory and neural responses across individuals

Given that intrinsic motivation was associated with memory and neural activity within individuals, we next asked whether these effects extended to shared memory and neural responses across individuals. Prior work has shown that greater attention to or engagement with a stimulus is associated with higher intersubject similarity in behavior and neural activity^48,49^, and that these shared responses predict subsequent memory^50–52^. To test whether wanting and liking similarly promote shared responses, we leveraged the passive-viewing condition. Unlike active seekers, who each encountered different content, viewers were exposed to the same content as one another and as the seeker who generated it, allowing direct comparisons across individuals.

We first examined whether events that both members of a viewer-viewer or viewer-seeker pair found more intrinsically motivating were more likely to be remembered by both (Fig. 7a). For each event and participant pair, pair-level wanting and liking were defined as the mean of the two participants’ ratings. Separate mixed-effects logistic regressions tested whether higher pair-level wanting or liking predicted shared recall. In viewer-viewer pairs, both pair-level wanting (*β* = 0.34, *z* = 16.21, *p* < 0.001, 95% CI = [0.30, 0.38]) and liking (*β* = 0.24, *z* = 11.34, *p* < 0.001, 95% CI = [0.20, 0.29]) were positively associated with shared recall. The same pattern was observed in seeker-viewer pairs, with both wanting (*β* = 0.41, *z* = 8.03, *p* < 0.001, 95% CI = [0.31, 0.51]) and liking (*β* = 0.27, *z* = 5.35, *p* < 0.001, 95% CI = [0.17, 0.37]) positively related to shared recall. These pair-level associations may largely reflect the within-person relationship between intrinsic motivation and memory reported above.

**Fig. 7.**
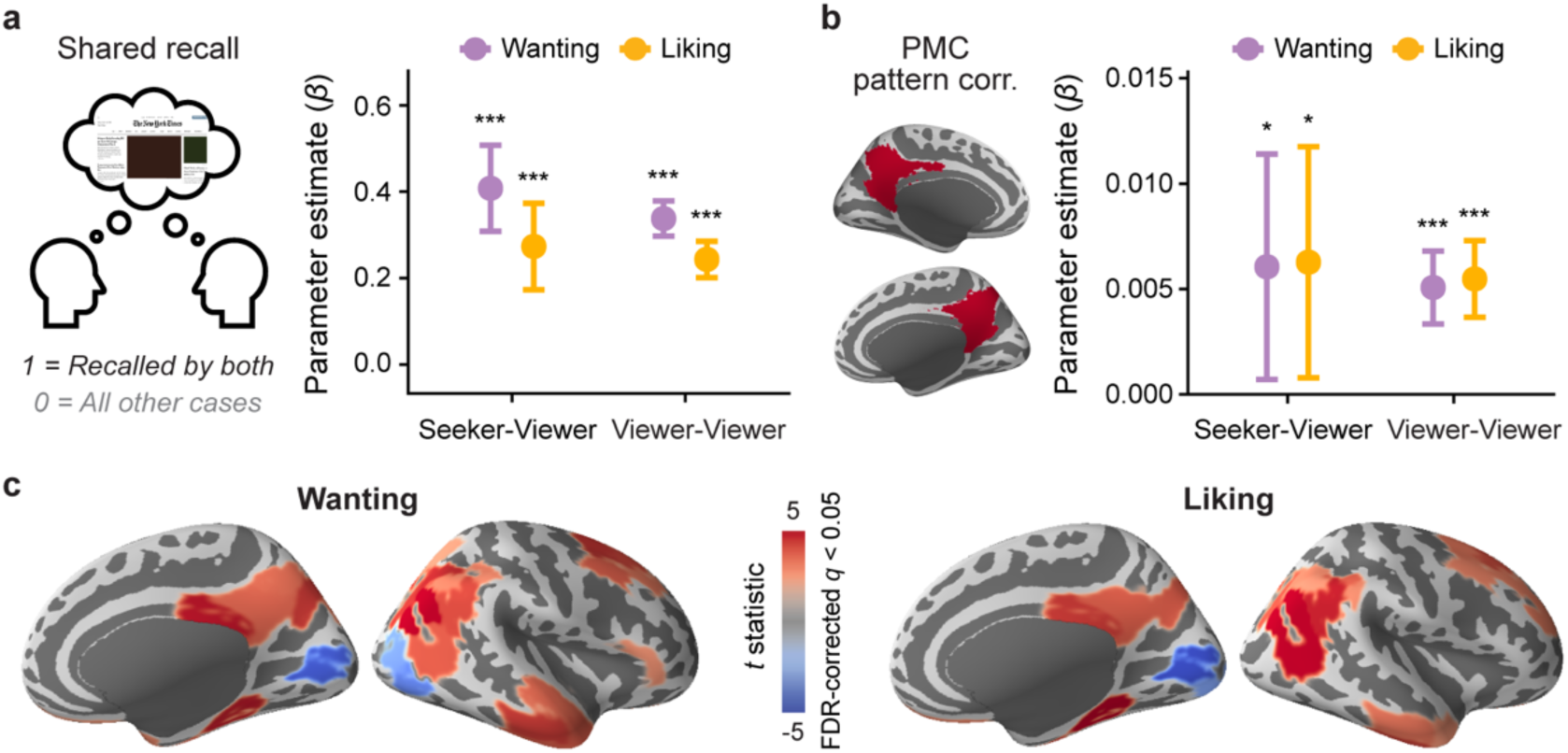
Effects of intrinsic motivation on shared memory and neural responses across individuals. (**a**) Effects of wanting (purple) and liking (yellow) on shared recall success in seeker-viewer and viewer-viewer pairs. Colored circles represent parameter estimates from logistic mixed-effects models predicting shared recall success (1 = both individuals in a pair recalled the event; 0 = at least one individual failed to recall the event) from wanting or liking ratings averaged across the two individuals in each pair. (**b**) Effects of wanting (purple) and liking (yellow) on event-wise intersubject pattern correlation in the posterior medial cortex (PMC; shown in red) during encoding (i.e., web browsing or screen recording video watching). Colored circles represent parameter estimates from linear mixed-effects models predicting PMC pattern similarity from wanting or liking ratings averaged across individuals within each seeker-viewer or viewer-viewer pair. In **a** and **b**, error bars indicate 95% confidence intervals. \**p* < 0.05, \*\*\**p* < 0.001. (**c**) Cortical regions in which event-wise viewer-viewer intersubject pattern correlation during encoding was associated with the pair’s mean wanting (left panel) or liking (right panel) ratings. Red and blue regions indicate positive and negative associations, respectively, as estimated using linear mixed-effects models. Statistical maps of *t* values are thresholded at *q* < 0.05 (FDR-corrected across all cortical parcels) and visualized on inflated medial (left) and lateral (right) surfaces of the right hemisphere. Full statistics for individual parcels are provided in Supplementary Tables 17-18.

We further tested whether pair-level intrinsic motivation was associated with shared neural representations of browsing events during information acquisition, quantified as event-wise intersubject pattern similarity. We first examined the posterior medial cortex (PMC), which showed event-specific neural reactivation in the previous analysis and is thought to represent high-level situation models^45^. Separate linear mixed-effects models (Fig. 7b) showed that higher pair-mean wanting (*β* = 0.005, *t* = 5.74, *p* < 0.001, 95% CI = [0.003, 0.007]) and liking (*β* = 0.005, *t* = 5.90, *p* < 0.001, 95% CI = [0.004, 0.007]) were each related to greater intersubject pattern similarity in PMC in viewer-viewer pairs. We found the same pattern in seeker-viewer pairs, with both wanting (*β* = 0.006, *t* = 2.22, *p* = 0.026, 95% CI = [0.001, 0.011]) and liking (*β* = 0.006, *t* = 2.25, *p* = 0.025, 95% CI = [0.001, 0.012]) positively related to PMC pattern similarity. An exploratory whole-brain analysis further showed that higher pair-level motivation in viewer-viewer pairs was associated with greater intersubject pattern similarity in regions within or near the default mode network^45,46^, primarily the PMC, lateral parietal cortex, and temporal cortex (Fig. 7c, Supplementary Tables 17-18). Seeker-viewer pairs showed similar positive relationships in lateral occipital-parietal regions (Supplementary Fig. 5). Thus, content that both members of a pair found more intrinsically motivating was more likely to be remembered by both and to elicit more similar neural responses.

## DISCUSSION

The current study investigated the neurocognitive processes associated with two independent dimensions of intrinsic motivation, wanting and liking, during naturalistic information seeking and learning. By combining a continuous, self-guided web-browsing task with functional neuroimaging and retrospective reports of moment-to-moment motivational states, we captured behavior and neural activity during active engagement with real web content, and related these processes to fluctuations in wanting and liking. Overall, the two motivational dimensions showed partially dissociable effects. Both predicted more stable information-seeking behavior but engaged overlapping yet distinct neural systems: wanting was more strongly associated with goal-directed cognition, whereas liking was linked to affective processing. Liking consistently predicted better memory, while wanting was beneficial primarily when liking was low. During passive viewing, wanting and liking were lower and more interdependent but still predicted memory and intersubject neural similarity.

Our naturalistic web browsing task let us characterize information-seeking behavior fully controlled by participants, including what content to seek from an almost unlimited range of options, and whether and when to move to new content. Higher wanting and liking predicted subsequent transitions to semantically related content and were generally accompanied by fewer content-switching actions, indicating more sustained engagement with the current content. These results suggest that intrinsically rewarding information reinforces prolonged consumption and further exploration of the same or related information, consistent with prior findings from controlled tasks^47,53^. Notably, liking showed a stronger negative relationship with content-switching actions than wanting, potentially reflecting its closer link to appreciation of the current stimulus^25^. A prior study using a virtual room exploration task reported a comparable dissociation between pre-exploration curiosity and post-exploration interest (conceptually analogous to wanting and liking, respectively): curiosity was linked to spatial exploratory actions, whereas interest tracked visual scanning but not spatial exploration^29^. Unlike that study, however, our task did not temporally separate pre– and post-exposure motivation, given the continuous nature of naturalistic browsing. Any effects of liking in our study may therefore reflect anticipation of pleasant information, affective responses to encountering it, or both.

During naturalistic web browsing, wanting and liking were associated with partially dissociable patterns of neural activity, supporting the view that they are distinct motivational dimensions^24,25^. Their neural correlates were widespread, extending beyond the reward system traditionally emphasized in information-seeking research^3,54^. First, higher wanting was related to greater activity in the lateral prefrontal cortex and caudate, consistent with curiosity-related activation in trivia paradigms^4,5^. These regions have been implicated in appraising the value of information and guiding goal-directed information seeking^7,8^. Greater activation at high wanting may therefore reflect higher perceived value of the information and greater cognitive control demands for planning how to close the information gap. In contrast, higher liking was related to lower activation in the anterior cingulate cortex and insula. Because these regions are commonly linked to cognitive conflict, interoceptive tension, and negative emotion^14,39,55^, reduced activation at higher liking may reflect a more positive affective state during information consumption.

Wanting and liking also engaged shared regions, as expected given their positive correlation. These included the superior and middle temporal cortices, hippocampus, and amygdala. The temporal regions support reading, language comprehension, and semantic integration^38,56^, which are central to web browsing, where participants continuously extract meaning from on-screen text. Greater activation in these regions is therefore consistent with deeper engagement with intrinsically motivating content. The positive association of hippocampal and amygdala activation with intrinsic motivation could reflect these regions’ role in novelty detection^57,58^, as novel or unpredicted information can induce curiosity and exploration^57^. These regions also showed wanting-by-liking interactions: the relationship between activation and wanting tended to be positive when liking was low but numerically negative when liking was high. One possible explanation is prediction error, often operationalized as the difference between pre-exposure curiosity and post-exposure satisfaction or interest^10,30,47^. Content initially low in wanting but unexpectedly pleasant could capture attention and promote further processing^59^, eliciting activation in novelty-detection and content-processing regions comparable to or greater than that for content high in both wanting and liking. Finally, both dimensions, especially liking, were generally negatively associated with activation in parts of the somatomotor and dorsal attention networks, including the frontal eye fields. We speculate that this is due to fewer browsing actions during highly motivating events. Although we controlled for actions in the analyses, residual activity related to higher-order motor processes or planned actions, such as eye movements, could remain. Future studies using tasks that require active behavioral control could more precisely measure overt actions, and even unexecuted movement plans, and separate them from motivation-related neural activity.

Intrinsic motivation during naturalistic web browsing shaped episodic memory for real-world online experiences. Both wanting and liking were linked to a higher probability of recall, with higher ratings for subsequently remembered than forgotten events. This aligns with the known benefits of curiosity and interest for learning and memory^2,8,41^. However, when their unique contributions were examined, only liking remained a significant predictor of recall. A small number of studies directly comparing pre-exposure curiosity and post-exposure interest have likewise reported stronger effects of post-exposure interest on memory for trivia answers^12,31^. Our study further showed that the more reliable contribution of liking extended to a neural marker of recall quality: encoding-retrieval pattern similarity in higher-order cortical regions thought to represent the content of specific events^43,44^. These results may reflect the mnemonic benefit of low-wanting, high-liking states, which produced the interaction in which wanting predicted recall only at low liking. This pattern mirrored the interaction in neural activation during browsing in a subset of regions positively associated with intrinsic motivation, including the hippocampus. As with neural activation, a plausible explanation for the memory pattern is that content eliciting high liking despite low wanting may generate positive prediction errors. Prior work has linked information-related prediction errors to enhanced memory^10,30,31^, paralleling the role of reward prediction errors in learning^60^.

Comparing the active information-seeking and passive-viewing groups allowed us to examine how agency, a core feature of real-world information seeking and learning, shapes intrinsic motivation and memory in a naturalistic setting. First, motivation ratings, especially wanting, were higher among seekers than viewers, consistent with prior findings that agency and autonomy enhance curiosity and interest^61,62^. With control over their actions, seekers could select personally motivating content, and the experience of choice itself may have been rewarding independent of content^63^. Second, wanting and liking were less strongly correlated among seekers, yielding a wider range of wanting-liking combinations. We speculate that this reflects seekers’ tendency to select self-relevant or personally useful information, such as work-or study-related materials, that they strongly wanted but did not necessarily find pleasant. Viewers, who had no personal stake in the selected content, may have rated such materials low in both wanting and liking, producing a tighter coupling between the two. This also suggest that self-directed information seeking is a particularly useful context for studying wanting and liking as distinct dimensions, including their unique contributions and interactions. Finally, despite these effects on motivation, recall performance did not differ consistently between seekers and viewers. Prior work on agency and memory is mixed: many studies report mnemonic benefits^64–66^, whereas others find benefits limited to particular memory types or no overall improvement^67,68^. Future research could clarify the conditions under which agency influences memory formation during naturalistic information seeking.

The passive-viewing condition, in which participants watched the same web content, also let us examine how intrinsic motivation relates not only to memory within individuals but also to memory shared across individuals. Events that both members of a pair rated higher in wanting or liking were more likely to be recalled by both, likely reflecting the generally positive relationship between intrinsic motivation and recall within individuals. Higher pair-level wanting or liking was also related to greater intersubject similarity in brain activation patterns, a neural marker previously shown to predict better subsequent memory^51,52^. Although prior studies have shown that emotional stimuli increase intersubject synchronization of neural responses^69,70^, little research has addressed the role of motivation for information seeking. One recent study found that participants more similar in the curiosity they felt while watching magic tricks also showed greater temporal alignment of neural activity in distributed cortical regions, including sensory, motor, and higher associative areas^17^. We extend these findings to intersubject similarity in the spatial distribution of activation, particularly in regions within and around the default mode network^45,46^, including the lateral and medial parietal cortices. Because activation patterns in these regions are sensitive to the semantic content of experiences^71^ and thought to represent internal models of ongoing events^45,46^, these results suggest that events higher in intrinsic motivation were represented or interpreted more consistently across individuals. One possibility is that higher wanting or liking kept participants focused on the current stimulus, whereas uninteresting material prompted stimulus-unrelated, idiosyncratic thoughts^72^, producing more diverse internal states. Alternatively, the causal direction may be reversed: content that is inherently ambiguous or hard to interpret for most people may elicit lower motivation.

Our study is subject to several methodological limitations. First, although self-guided web browsing is an ecologically valid task rarely used in neuroimaging, participants’ browsing may not have matched their everyday internet use. Awareness of the experimenters and the screen recording may have led participants to select more socially acceptable content. Prohibiting logins to personal social media accounts, for privacy, is a notable limitation, given the prevalence of social media use and its potential impact on mental health and brain activity^73^. Future studies could examine brain activity during even more realistic information seeking by allowing more unrestricted browsing. Second, we collected retrospective ratings of wanting and liking to avoid disrupting the natural flow of browsing, but reliance on memory for these ratings may have introduced noise or bias from forgetting or memory distortion. Ratings were also limited to 6-second resolution to prevent participant fatigue during the task. Future work could employ finer-grained windows to better capture fluctuations in motivation. Third, wanting and liking ratings were naturally correlated, consistent with prior reports^12,47^. Although we still observed dissociations between them in several respects, this correlation may have limited the range of distinct motivational states available for analysis. For example, the high correlation between wanting and liking during passive viewing prevented us from examining their interaction in that condition. Finally, our study does not address why specific content elicited higher or lower intrinsic motivation. Prior work suggests that information consistent with existing beliefs^74^ or facilitating efficient knowledge structures^75^ may be more valued. External contextual factors and individual differences in personality and life history may also shape motivation for specific information^20^. Further research is needed to determine whether and how wanting and liking for diverse and complex real-world content can be predicted across individuals.

Overall, using a self-guided web-browsing fMRI paradigm, we found that wanting and liking, two theoretically proposed independent dimensions of intrinsic motivation^24,25^, make both shared and distinct contributions to how the human brain seeks, represents, and remembers information in naturalistic settings. Beyond informing perspectives on the motives underlying information seeking, our study extends prior behavioral and neural research on motivated learning^41^ to a more ecologically valid context in which individuals freely and continuously navigate an expansive knowledge space. This approach may promote future investigation of the neurocognitive mechanisms underlying the cycle of real-world information seeking and learning, whereby knowledge acquisition generates new questions that drive further exploration^2^. The web-browsing fMRI paradigm may also facilitate research into the neural mechanisms of diverse naturalistic online behaviors associated with mental health conditions such as addiction and depression^34,73,76^, as well as societal phenomena such as political polarization^77^.

## METHODS

### Active information seeking participants

Fifty-six healthy adults (30 females; mean age = 23.14 years, range: 18-35 years) were recruited from Purdue University and the Greater Lafayette community. All participants reported no history of neurological or psychiatric disorders and had normal hearing and normal or corrected-to-normal vision. Before participation, they completed an MRI safety screening and provided written informed consent in accordance with protocols approved by the Purdue University Institutional Review Board (approval number: IRB-2023-1089). Participants received monetary compensation for their participation.

Participant exclusions were performed separately for different analyses to maximize the available sample size for each analysis. The final behavioral sample included 54 participants (30 females; mean age = 23.26 years) after excluding two participants: one because of difficulty controlling the MR-compatible mouse and reading text on the screen, and the other because they did not complete the post-scan ratings. The web browsing fMRI analyses included 53 participants (29 females; mean age = 23.32 years) after excluding one additional participant because of MRI signal artifacts. The narrated recall fMRI analyses included 48 participants (25 females; mean age = 23.38 years) after excluding five additional participants because of excessive head motion during the recall task (mean framewise displacement > 0.5 mm). Additional analysis-specific exclusions were applied where appropriate. For example, one participant had no variance in wanting ratings and was excluded from analyses involving wanting. Another participant had no variance in keypress counts (all values were zero) and was excluded from analyses that included keypress counts.

### Active information seeking experimental design and procedures

Participants completed an fMRI session involving naturalistic web browsing followed by a narrated free recall task (Figs.1a-b). After scanning, they rated their subjective motivation and completed questionnaires (Fig. 1c).

#### Web browsing task

Participants freely explored real-world web content during two 15-minute fMRI runs. Each run began with the Google Chrome start page, which included a search bar and 10 shortcut links (e.g., Wikipedia, YouTube, and The New York Times). The shortcuts were selected from a list of the most visited websites in the United States (https://www.similarweb.com/top-websites, December 2023), excluding websites that required login to personal accounts and pornographic websites. Browsing began upon presentation of a pop-up prompt (“Begin web browsing!”), which was time-locked to the acquisition of the first fMRI volume.

Participants used an MR-compatible mouse (Nata Technologies Inc.) and an on-screen keyboard to browse and search the web as they normally would in daily life, subject to a few constraints. They were encouraged to visit multiple websites rather than spending extended periods on a single website or watching any one video for more than approximately 3-4 minutes. To protect participant privacy, logging in to personal email or social media accounts was prohibited. To maintain experimental consistency, participants were also discouraged from visiting non-English websites or playing online games. At the end of the first run, participants stopped browsing when the scanner stopped and waited for the second run to begin. When the “Begin web browsing!” prompt reappeared, they resumed browsing from where they had left off.

Participants’ browsing activity was screen-recorded throughout the task. Manual information-seeking actions were continuously recorded using the pynput Python library (https://pypi.org/project/pynput/). Specifically, mouse clicks, scrolls, and keypresses were recorded and converted into volume-wise binary variables, with volumes containing a given action coded as 1 and all other volumes coded as 0. Mouse movement was quantified by recording cursor positions and calculating the distance traveled every 0.1 s. The resulting distances were summed within each 1.5-s interval to match the fMRI repetition time (TR).

In this and all other fMRI runs, visual stimuli were presented using a Hyperion HD (1080p) MRI rear-projection system, which projected images onto a screen positioned behind the magnet bore. Participants viewed the screen through an angled mirror mounted on the head coil. Audio was delivered through in-ear headphones (Avotec Silent Scan 3300).

#### Free recall task

Following the web browsing task and subsequent anatomical and field map scans, participants completed a narrated free recall task during fMRI. They were instructed to verbally describe everything they remembered from the browsing session in as much detail as possible, including their thoughts and feelings. The task began with on-screen instructions, followed by a brief “Wait” message while the experimenter prepared the scanner. The word “Begin!” then appeared on the screen for 1.5 s, prompting participants to start speaking. All instructions and messages were presented in white text on a gray background. A small central dot remained on the screen throughout the task, although participants were not required to fixate on it. Participants were instructed to speak continuously for at least 10 min and to say “I’m done” when they had finished recalling. To minimize motion artifacts, participants were instructed to minimize jaw movement while speaking and to avoid moving their heads. Participants’ verbal responses were recorded using a MR-compatible recording system (FOMRI II; Optoacoustics Ltd.) and Audacity software (https://www.audacityteam.org).

#### Subjective ratings and surveys

After the MRI scan, participants provided retrospective moment-to-moment ratings of their intrinsic motivation during web browsing while watching screen recordings of their own browsing activity at double speed. The video was presented in successive 3-s playback segments, each corresponding to 6 s of the original browsing recording. After each segment, playback paused, and participants rated their wanting and liking of the just-viewed content on a 5-point Likert scale. The wanting prompt, “How strongly did you want to know the information you were browsing for?”, appeared first, followed by the liking prompt, “How strongly did you like the information you were browsing for?”. Both questions were presented in white text on a gray background, and participants responded at their own pace. Playback then resumed from the point at which it had paused.

After rating their intrinsic motivation, participants completed self-report questionnaires assessing demographic characteristics, mental health, and personality. These included the Big Five Inventory^78^, the Mind Wandering Questionnaire^79^, the Internet Addiction Test^80^, the Adult ADHD Self-Report Scale^81^, the Depression Anxiety Stress Scales-2^82^, and the Curiosity and Exploration Inventory^83^. Participants also answered three study-specific questions assessing whether they visited websites they normally use during the experiment, how frequently they typically visit those websites in daily life, and whether the recommended websites led them to visit websites they do not usually visit.

### Passive viewing participants

A total of 25 healthy adults were recruited for the passive-viewing condition independently of the sample recruited for the active information-seeking condition. Participants were randomly assigned to watch one of two selected screen recordings of web browsing performed by participants from the active information-seeking condition (Video 1 or Video 2). All participants reported no history of neurological or psychiatric disorders, had normal hearing and normal or corrected-to-normal vision, and passed an MRI safety screening. They provided written informed consent in accordance with protocols approved by the Purdue University Institutional Review Board and received monetary compensation for their participation.

Two participants were excluded from the analyses: one because they fell asleep during the experiment and the other because of excessive head motion throughout the scan (mean framewise displacement > 0.5 mm). The final analytical sample included 23 participants (13 females, 9 males, and 1 nonbinary; mean age = 20.8 years, range = 18-42 years), who did not significantly differ from the active information-seeking sample on any of the survey measures collected (Supplementary Table 13). Of these participants, 11 watched Video 1 and 12 watched Video 2. The two video groups also did not significantly differ on any of the survey measures (Supplementary Table 14).

### Passive viewing experimental design and procedures

Participants completed an experimental session consisting of a video watching task and a free recall task during fMRI, followed by post-scan ratings and survey assessments (Fig. 1).

#### Video watching task

Instead of browsing the web themselves, participants watched one of two screen recordings of web browsing sessions performed by two participants from the active information-seeking condition. Because data collection for the passive-viewing condition began while recruitment for the active information-seeking condition was still ongoing, the stimuli were selected from among the first 30 participants in the active information-seeking condition. The two screen recordings were chosen based on several data quality criteria, including the absence of technical or procedural issues throughout the experiment, minimal head motion during scanning, and relatively long and detailed recall. The video-watching phase consisted of two functional runs. Each run presented one 15-min half of the recorded web browsing session, preserving the original order of the browsing activity. Participants were instructed to attend to the videos and their own subjective feelings to the viewed content. No verbal or manual responses were required during this phase.

#### Free recall task

Participants were instructed to verbally recall everything they remembered from the video-watching phase, including their thoughts and feelings, using the same procedures as in the active information-seeking condition.

#### Subjective ratings and surveys

After the MRI scan, participants provided retrospective moment-to-moment ratings of their intrinsic motivation during the video-watching phase. The rating procedure was identical to that used in the active information-seeking condition, except that the wanting (“How strongly did you want to know the information you were watching?”) and liking (“How strongly did you like the information you were watching?”) prompts were modified to reflect the passive viewing task. The post-scan surveys were also identical to those used in the active information-seeking condition, except that the three questions about participants’ website use were omitted.

### Behavioral data and stimuli preprocessing

#### Event segmentation and timestamping of web browsing screen recording

Ten human annotators manually segmented each participant’s web browsing screen recordings into a nested hierarchical structure consisting of three levels of granularity: websites, webpages, and fine-grained events. Each annotator processed an average of 5.4 participants’ screen recordings (range = 1-20).

Annotators first identified the website based on the domain name displayed in the browser’s URL bar (e.g., google.com) and labeled each webpage using the title shown in the browser tab (e.g., MRI –Google Search). Within each webpage, they further segmented the screen recording into fine-grained events, each corresponding to a distinct information-seeking action, such as scrolling through search results, reading comments, closing a pop-up advertisement, and watching a video. Actions lasting several tens of seconds or longer, such as reading a long article or watching a lengthy YouTube video, were further divided into multiple fine-grained events. For example, different scenes within a YouTube video were treated as separate events, with event boundaries defined by changes in features such as topic, location, action, or characters. When participants continued browsing after the end of the 15-minute scanning run, events occurring beyond the scan endpoint were not annotated. Finally, annotators recorded the onset time of each fine-grained event with frame-level precision. Because adjacent events were temporally contiguous in the screen recordings, end times were not recorded separately.

In addition to the initial three-level segmentation, a mid-grained level of segmentation was introduced subsequently. Because fine-grained events were defined based on individual actions, adjacent events often contained highly similar webpage content. For example, scrolling down and then back up on the same webpage was annotated as two separate fine-grained events despite involving essentially the same content. Moreover, many fine-grained events were relatively brief for fMRI analyses (mean duration = 8.32 s, SD = 3.47 s) given the hemodynamic response delay. To address these issues, a senior annotator (C.K.) grouped semantically related, temporally consecutive fine-grained events within the same webpage into mid-grained events. Each mid-grained event represented a continuous episode of engagement with a single piece of content, such as an article, video, or social media post, and served as the primary unit of analysis in the current study. The senior annotator also reviewed all screen recordings to ensure the quality and consistency of the event segmentation.

#### Annotation of web browsing events

Seven human annotators generated text descriptions for each mid-grained event from the screen recordings of active information-seeking participants included in the analyses. Each annotator processed an average of 7.71 screen recordings (range = 1-24). Annotators watched each screen recording and wrote a one-to three-sentence summary describing the corresponding mid-grained event. Summaries were intended to capture the primary content the participant was attending to during each event. Annotators were instructed to focus on webpage content rather than the participant’s actions, except when the action itself constituted the primary content (e.g., typing search terms using the on-screen keyboard).

#### Audio transcription and timestamping of narrated recall

Each participant’s narrated free recall audio recording was manually transcribed and timestamped by human annotators. A total of 12 annotators processed 77 recordings included in the analyses across the active information-seeking and passive-viewing conditions. Each annotator processed an average of 6.42 recordings (range = 1-28). Recordings were first transcribed verbatim, including fillers such as “uh” and “um.” The transcripts were then segmented into individual sentences, and long sentences containing multiple clauses were further divided into separate clauses. Annotators subsequently recorded the onset and offset times of each segment with millisecond resolution while listening to the audio recordings and visually inspecting the corresponding waveforms and spectrograms. Recall segments that ended before the scan began or began after the scan ended were excluded from analysis.

#### Alignment of narrated recall to web browsing events

To identify which web browsing events (performed or observed) were successfully remembered, 11 human annotators manually aligned each participant’s recall segments with the annotated events from the corresponding screen recording (Fig. 4a). Each annotator processed an average of 7 participants’ recall transcripts (range = 1-25) across the active information-seeking and passive-viewing conditions. Annotators first watched the screen recording and listened to the participant’s recall recording to familiarize themselves with the browsing session and the recall content. For each recall segment (a sentence or clause), annotators identified all fine-grained event numbers corresponding to the recalled content. At the mid-grained event level, an event was classified as remembered if at least one of its constituent fine-grained events was matched to the participant’s recall; otherwise, it was classified as forgotten. This remembered/forgotten classification served as the binary memory outcome in subsequent analyses. Annotators also identified segments consisting of filler utterances, task-irrelevant utterances (e.g., “It’s cold in here, by the way.”), or task-relevant utterances that were too general to be aligned with specific events (e.g., “I browsed a lot of different websites.”). These segments were excluded from analysis. Finally, 6 annotators, 4 of whom had also participated in the initial recall-video event alignment, independently reviewed the transcripts they had not originally annotated, and corrected errors such as typographical mistakes (mean = 6.49 transcripts per annotator; range = 5-20).

#### Extracting screen recording brightness and audio amplitude

To quantify low-level sensory characteristics of the web browsing screen recordings, we extracted two time-resolved stimulus features: screen brightness and audio amplitude. Both features were computed in temporal windows aligned with the fMRI TR. For screen brightness, each video was read frame by frame from the onset of the experiment, and each frame was converted to grayscale. The mean pixel intensity of each grayscale frame was calculated and then averaged within consecutive, non-overlapping 1.5-s TR windows to obtain TR-wise brightness values. For audio amplitude, the audio track was extracted from each screen recording, downsampled to 8000 Hz, and converted to a single channel. Audio amplitude was quantified as the mean absolute waveform amplitude within consecutive, non-overlapping 1.5-s TR windows from experimental onset, yielding TR-wise audio amplitude values. The TR-wise screen brightness and audio amplitude measures were subsequently averaged across all TRs corresponding to each mid-grained web browsing event to generate event-level feature values. These event-wise features were included as regressors in linear mixed-effects models to account for the contributions of low-level perceptual features to brain responses during naturalistic information seeking.

### Behavioral data analysis

#### Web-browsing actions

To test whether intrinsic motivation was associated with information-seeking actions during naturalistic web browsing, we computed event-wise mean values for browsing actions (clicks, scrolls, keypresses, mouse movement distance) and motivation ratings (wanting, liking) by averaging their TR-wise values within each event (Fig. 2b). For each participant, Pearson correlation coefficients were then calculated between event-wise motivation ratings and each browsing action. The resulting correlation coefficients were Fisher *z*-transformed and tested against zero using two-tailed one-sample *t*-tests. Two-tailed paired-samples *t*-tests were used to compare the motivation-action correlations across wanting and liking. Additionally, for exploratory purposes, we computed correlations among all pairs of motivation ratings, browsing actions, and low-level perceptual features of the screen recording videos (Supplementary Fig. 1).

#### Event-to-event semantic continuity during web browsing

To test whether intrinsic motivation was associated with the semantic continuity of information-seeking trajectories, we quantified the semantic similarity between each event and the event that immediately followed it (Fig. 2b). The human-annotated text description of each event was embedded into a 768-dimensional vector using the pretrained Sentence-BERT model (all-mpnet-base-v2) implemented in the Sentence Transformers Python package (v2.2.0; https://www.sbert.net). Semantic similarity for each consecutive event pair was computed as the cosine similarity between their embedding vectors. For each participant, Pearson correlation coefficients were calculated between event-wise motivation ratings and event-to-event semantic continuity. The resulting correlation coefficients were Fisher *z*-transformed and tested against zero using two-tailed one-sample *t*-tests. A two-tailed paired-samples *t*-test was used to compare the correlations between wanting and liking.

#### Recall performance in active information seeking

To examine whether intrinsic motivation influences memory performance, we compared motivation ratings between subsequently remembered and forgotten events. For each participant, mean wanting and liking ratings were computed separately for remembered and forgotten events. Differences in motivation between remembered and forgotten events were then tested across participants using two-tailed paired-samples *t*-tests.

We next examined how wanting and liking interacted to influence subsequent memory while accounting for potential confounds. We fit a mixed-effects logistic regression with the binary memory outcome (remembered = 1, forgotten = 0) as the dependent variable. Fixed effects included the main effects of wanting, liking, and their interaction. Covariates included the mean values of participant actions per event (clicks, scrolls, keypresses, and mouse movement distance), run number (first or second run), and low-level perceptual features (mean screen brightness and audio amplitude). Participant was included as a random intercept. Wanting and liking ratings were centered at the midpoint of the 5-point Likert scale. Participant actions and low-level perceptual features were z-scored within participants. The model was specified as: recall success ∼ wanting + liking + wanting × liking + run + screen brightness + audio amplitude + click + scroll + keypress + mouse movement distance + (1 | participant).

#### Recall performance in passive viewing

To examine the association between intrinsic motivation and event-level recall success in the passive-viewing condition, we fit mixed-effects logistic regression models analogous to those used for the active information-seeking condition. Because wanting and liking ratings were highly correlated during passive viewing (mean *r* = 0.82, SD = 0.14), they were entered into separate models, and their interaction was not considered. Each model included either wanting or liking as the predictor of interest, with video stimulus (Video 1 or Video 2), scanning run (first or second), screen brightness, and audio amplitude included as fixed-effect covariates. Because participants in the passive-viewing condition (viewers) did not make any behavioral responses during video watching, browsing action covariates were not included. The model was specified as: recall success (1 = remembered, 0 = forgotten) ∼ motivation (wanting or liking) + video + run + screen brightness + audio amplitude + (1 | viewer).

#### Shared recall success across individuals

We fit mixed-effects logistic regression models to examine whether intrinsic motivation during encoding was associated with successful recall shared across individuals. Shared recall was defined as whether both members of a pair successfully remembered a given event (1 = remembered by both individuals, 0 = otherwise). First, we examined viewer-viewer pairs by testing whether shared recall was predicted by the pair’s mean wanting or liking rating for each event. All possible pairs of viewers who watched the same screen recording video were included, with pair identity included as a random intercept. Separate models were fit for wanting and liking. The model was specified as: shared recall ∼ viewer-viewer mean motivation (wanting or liking) + video + run + screen brightness + audio amplitude + (1 | pair).

Following the same approach, we next tested whether pair-level wanting or liking predicted shared recall between each viewer and the participant in the active information-seeking condition (the seeker) whose screen recording the viewer watched. Again, all possible seeker-viewer pairs were included. Because seeker identity was completely determined by the video variable included as a fixed effect, viewer identity, rather than pair identity, was included as a random intercept. Separate mixed-effects logistic regression models were fit for wanting and liking. The model was specified as: shared recall ∼ seeker-viewer mean motivation (wanting or liking) + video + run + screen brightness + audio amplitude + (1 | viewer).

#### Comparing motivation and memory between active seeking and passive viewing

We compared the active information-seeking and passive-viewing conditions in terms of mean wanting and liking ratings, the similarity between wanting and liking, and memory performance. For each participant, mean wanting and liking were calculated by averaging event-wise ratings across all events. The similarity between wanting and liking was quantified as the Fisher *z*-transformed Pearson correlation between event-wise wanting and liking ratings within each participant. Memory performance was defined as the proportion of events successfully recalled.

For each behavioral measure, we performed two complementary comparisons. First, we compared the active information-seeking and passive-viewing groups. Separately for each video presented in the passive-viewing condition, viewers who watched that video were compared with all participants in the active information-seeking condition using two-tailed Welch’s independent-samples *t*-tests. Second, we compared viewers with the corresponding seeker who generated the screen recording they watched. Specifically, separately for each video, viewers were compared with the seeker using two-tailed one-sample *t*-tests.

### Neuroimaging data acquisition and analysis

#### Neuroimaging data acquisition

MRI data were collected using a Siemens 3T scanner at the Purdue Life Science MRI facility. Functional images were acquired with a T2*-weighted multiband echo-planar imaging (EPI) sequence (TR = 1.5 s, TE = 30 ms, flip angle = 52°, multiband factor = 4, 60 oblique axial slices, matrix = 110 × 106, voxel size = 2 × 2 × 2 mm³). B0 field inhomogeneities were collected using a field map sequence with the same geometry. High-resolution structural images were acquired using a T1-weighted MPRAGE sequence (TR = 2.3 s, TE = 2.98 ms, flip angle = 9°,176 axial slices, matrix = 248 × 256, voxel size = 1 × 1 × 1 mm³).

#### Neuroimaging data preprocessing

MRI data were first organized into the Brain Imaging Data Structure (BIDS) format. Preprocessing of high-resolution anatomical images were performed using the recon-all pipeline of FreeSurfer (version 7.3.2). Functional images were preprocessed using fMRIprep (version 23.2.1) with default settings. Specifically, functional images were corrected for head motion and B0 magnetic inhomogeneity. Functional images were then coregistered to the anatomical image and resampled to the MNI152 volume space. Additionally, functional images were smoothed (FWHM 4 mm) in the volume space using nltools 0.5.0 (https://nltools.org/). Nuisance signals were regressed out using a linear regression model that included linear and quadratic trends, six head motion parameters (translations and rotations along x, y, and z axes), framewise displacement, and average signals from cerebrospinal fluid and white matter. The resulting time series were z-scored within each voxel across all volumes within the scanning run. Volumes with framewise displacement ≥ 1 mm were excluded from further analysis.

#### Cortical parcellation and region of interest (ROI) definition

For whole-brain cortical analyses, we used the functionally defined cortical parcellation atlas developed by Schaefer and colleagues^36^, which is based on resting-state functional connectivity. We used the version of the atlas that divides the cortical surface into 200 parcels (100 per hemisphere), organized into 17 large-scale functional networks identified in a previous study^84^. Using the atlas, we also defined a bilateral posterior medial cortex (PMC) ROI by merging seven precuneus/posterior cingulate cortex (pCunPCC) parcels from the Default Network: LH_Default_pCunPCC_1-4 and RH_Default_pCunPCC_1-3.

In addition to the cortical regions, we examined a set of anatomically defined a priori subcortical ROIs implicated in motivation and memory, including the hippocampus, amygdala, caudate, and putamen (Supplementary Fig. 3). Bilateral masks for these regions were derived from each participant’s FreeSurfer subcortical segmentation of the high-resolution anatomical image and transformed into MNI152 template space. To minimize partial-volume effects from adjacent tissue, each mask was eroded by one voxel (1 mm).

#### Brain activation associated with intrinsic motivation

To examine the association between intrinsic motivation and neural activation during naturalistic web browsing, we fit linear mixed-effects models predicting event-wise neural activation from wanting and liking ratings. For each participant and brain region, event-wise neural activation was computed by averaging preprocessed fMRI signals across voxels within the region and across TRs corresponding to each event. To account for the hemodynamic response delay, event windows were shifted forward by 4.5 s (3 TRs). Events at the end of each scanning run were excluded if all corresponding TRs were unavailable after the temporal shift.

For each brain region, we first fit two separate models, each including either wanting or liking as the predictor of interest. Scanning run, low-level perceptual features, and browsing actions were included as covariates. Participant was included as a random intercept. The model was specified as: neural activation ∼ motivation (wanting or liking) + run + screen brightness + audio amplitude + click + scroll + keypress + mouse movement distance + (1 | participant).

Next, we fit an additional model for each brain region that included wanting, liking, and their interaction as predictors. This allowed us to examine the unique effect of each motivational dimension after controlling for the other, as well as to test whether the two dimensions interacted to affect neural activation. The model was specified as: neural activation ∼ wanting + liking + wanting × liking + run + screen brightness + audio amplitude + click + scroll + keypress + mouse movement distance + (1 | participant).

#### Brain activation associated with successful memory formation

We fit mixed-effects logistic regression models to identify brain regions whose activation during web browsing predicted successful memory formation. For each brain region, subsequent recall success (remembered = 1, forgotten = 0) was modeled as a function of event-wise neural activation, which was computed using the same procedure described above for the analysis of brain activation associated with intrinsic motivation. To estimate the unique contribution of neural activation, the models additionally included scanning run, low-level perceptual features, and browsing actions as fixed-effect covariates. Participant was included as a random intercept. The model was specified as: recall success ∼ neural activation + run + screen brightness + audio amplitude + click + scroll + keypress + mouse movement distance + (1 | participant).

#### Encoding-retrieval pattern similarity

To quantify neural reinstatement of web browsing events during recall, we computed event-specific encoding-retrieval pattern similarity (Fig. 5b). For each participant and brain region, event-specific multivoxel activity patterns were obtained by averaging preprocessed voxelwise fMRI signals across the TRs corresponding to each event during both web browsing (encoding) and narrated recall (retrieval). Event timestamps were shifted forward by 3 TRs (4.5 s), and events at the end of each scanning run were excluded if all corresponding TRs were unavailable after the temporal shift. For each recalled event, we calculated the Fisher *z*-transformed Pearson correlation between its retrieval activity pattern and the encoding activity pattern of every web browsing event. Correlations between matching encoding and retrieval events were classified as matched, whereas correlations between different events were classified as mismatched. Correlation values were then averaged separately for matched and mismatched pairs within each participant. Finally, matched and mismatched encoding-retrieval pattern similarity values were compared across participants using a two-tailed paired-samples *t*-test. Greater encoding-retrieval pattern similarity for matched than for mismatched event pairs was considered to reflect event-specific neural reinstatement of encoding-related activity during recall.

Next, for cortical parcels showing significant event-specific neural reinstatement, we fit linear mixed-effects models to examine whether reinstatement strength was associated with intrinsic motivation. Only encoding-retrieval similarity for matched pairs was considered as the dependent variable. Fixed effects included the main effects of wanting and liking and their interaction. Scanning run, browsing actions, and low-level perceptual features were included as covariates, and participant was included as a random intercept. The model was specified as: encoding-retrieval similarity ∼ wanting + liking + wanting × liking + run + screen brightness + audio amplitude + click + scroll + keypress + mouse movement distance + (1 | participant).

#### Event-specific intersubject neural pattern similarity

We performed intersubject pattern similarity analysis to test whether intrinsic motivation was associated with shared neural representations of information acquisition experiences across individuals. For each participant and brain region, event-level neural activation patterns during encoding were obtained using the same procedures described above for the encoding-retrieval pattern similarity analysis.

For each brain region, we first examined intersubject pattern similarity among viewers who watched the same screen recording video. For every possible viewer-viewer pair, event-specific intersubject pattern similarity was computed as the Fisher *z*-transformed Pearson correlation between the participants’ event-level encoding activation patterns for the same event. We then fit linear mixed-effects models to test whether event-wise intersubject pattern similarity was predicted by the pair’s mean wanting or liking rating for each event. Because wanting and liking were highly correlated during passive viewing, separate models were fit for the two motivational dimensions. Each model included either wanting or liking as the predictor of interest, with video, scanning run, and low-level perceptual features included as fixed-effect covariates. Viewer-viewer pair was included as a random intercept. The model was specified as: viewer-viewer intersubject pattern similarity ∼ viewer-viewer mean motivation (wanting or liking) + video + run + screen brightness + audio amplitude + (1 | pair).

Following the same approach, we next tested whether pair-level wanting or liking predicted intersubject pattern similarity between each viewer and the seeker whose screen recording the viewer watched, separately for each brain region. For every possible viewer-seeker pair, event-wise intersubject pattern similarity during encoding was computed and used as the dependent variable in linear mixed-effects models. Because seeker identity was completely determined by the video variable included as a fixed effect, viewer identity, rather than pair identity, was included as a random intercept. Separate models were fit for wanting and liking. The model was specified as: seeker-viewer intersubject pattern similarity ∼ seeker-viewer mean motivation (wanting or liking) + video + run + screen brightness + audio amplitude + (1 | viewer).

### Statistical tests

Details of the statistical procedures for each analysis are provided in the corresponding Materials and Methods subsections. For comparisons of means, we used Student’s *t*-tests, except when comparing seekers and viewers, for which we used Welch’s *t*-tests to account for unequal variances and the small sample size of each video viewer group. For within-participant comparisons, participants with missing data in any condition were excluded from analysis. All reported statistical tests were two-tailed. Multiple comparisons across variables or brain regions were corrected using the Benjamini-Hochberg false discovery rate procedure.

Mixed-effects model analyses were performed in R (version 4.4.1) using the lme4 package. Specifically, mixed-effects logistic regression models predicting binary memory outcomes (remembered vs. forgotten) were fit using the glmer function, while linear mixed-effects models predicting continuous outcomes, such as encoding-recall pattern similarity and neural activation measures, were fit using the lmer function. Across all mixed-effects models, wanting and liking ratings were centered at the midpoint of the scale by subtracting 3 from each rating (original scale: 1–5; centered range: −2 to 2). Continuous nuisance regressors, including browsing actions and low-level perceptual features, were *z*-scored within participants, and run was included as a categorical covariate. All models included a random intercept for participant. For linear mixed-effects models with continuous outcomes, fixed-effect coefficients were evaluated using *t*-tests with Satterthwaite-approximated degrees of freedom. For generalized linear mixed-effects models with binary outcomes, fixed-effect coefficients were evaluated using Wald *z*-tests.

For significant interaction terms, the nature of the interaction was examined using simple slopes analysis, implemented with the sim_slopes function in the interactions package. Specifically, the slope of the focal predictor was estimated at low and high levels (±1 SD from the mean) of the moderator variable. Simple slopes were tested using the inferential method corresponding to the fitted model: Satterthwaite-approximated *t*-tests for linear mixed-effects models and Wald *z*-tests for binomial generalized linear mixed-effects models.

## DATA AVAILABILITY

The raw neuroimaging data will be deposited on OpenNeuro.org upon publication of this study.

## CODE AVAILABILITY

All analyses in this study were conducted using publicly available MRI data processing software (e.g., FSL, FreeSurfer) and Python packages. The analysis scripts used to generate the display items in this paper will be made available on GitHub. Other analysis scripts will be available upon request from the corresponding authors (H.S. and H.L.).

## ACKNOWLEDGMENTS

We thank Morgan Bailey, Camille Deakins, Chenrui Yin, Yuna Lee, Alyssa McMahon, Jared Ernstes, Dylan Nunes, Brooke Burks, and Soroosh Kermani for assistance with data collection; Yoolim Hong, Gang Chen, Taylor Korn, Matthew Sindac, Carl Xu, Austin Lugo, Benjamin Dravis, and Caden Wright for assistance with data analysis. This research was supported by the Purdue University MRI Pilot Grant (H.L.) and the Indiana Clinical and Translational Sciences Institute Pilot Funding for Research Use of Core Facilities (H.L.)

## AUTHOR CONTRIBUTIONS

Conceptualization: H.S., H.L.

Methodology: H.S., H.L.

Investigation: H.S., C.K., P.R.L., H.L.

Visualization: H.S., H.L.

Supervision: H.L.

Writing—original draft: H.S., H.L.

Writing—review & editing: H.S., H.L.

