## Supplementary Materials for "Neurocognitive signatures of intrinsic motivation in naturalistic information seeking and memory"

### SUPPLEMENTARY FIGURES

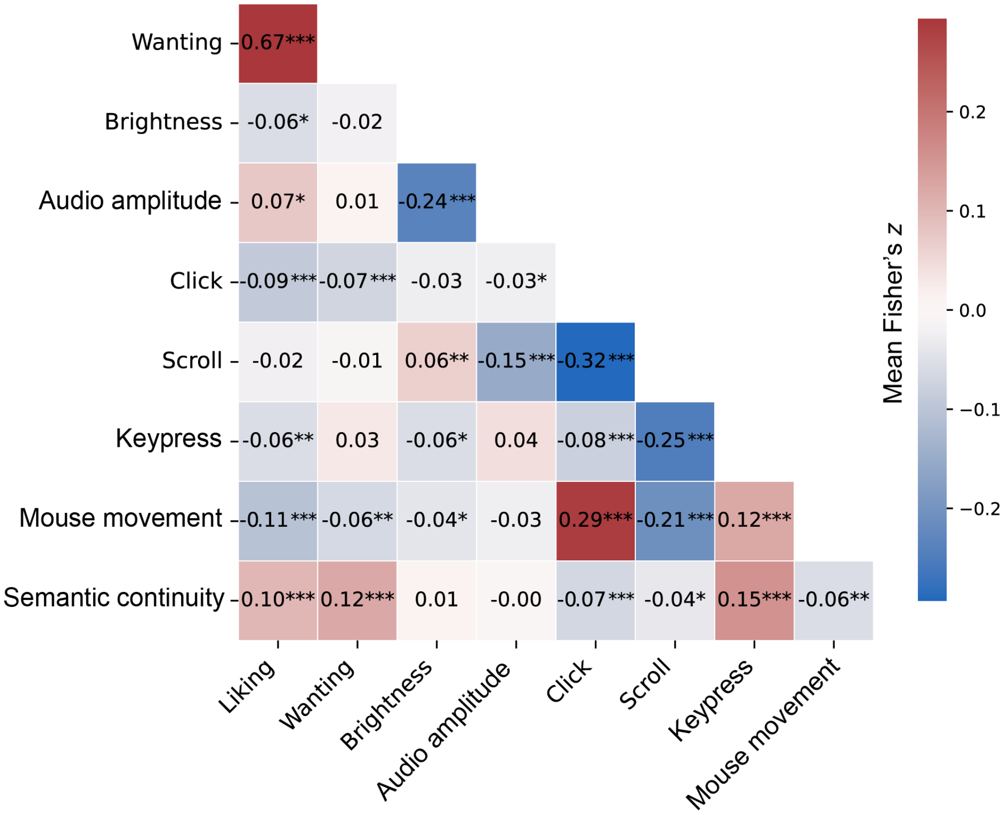


**Supplementary Fig. 1. Correlations (Fisher's *z*-transformed Pearson *r*) among motivation ratings, low-level audiovisual features, and browsing actions.** Correlations were computed within each participant and then averaged across participants. Warm colors indicate positive correlations, and cool colors indicate negative correlations. Statistical significance was assessed using two-tailed one-sample *t*-tests against zero. *$p < 0.05$, **$p < 0.01$, ***$p < 0.001$ (uncorrected)

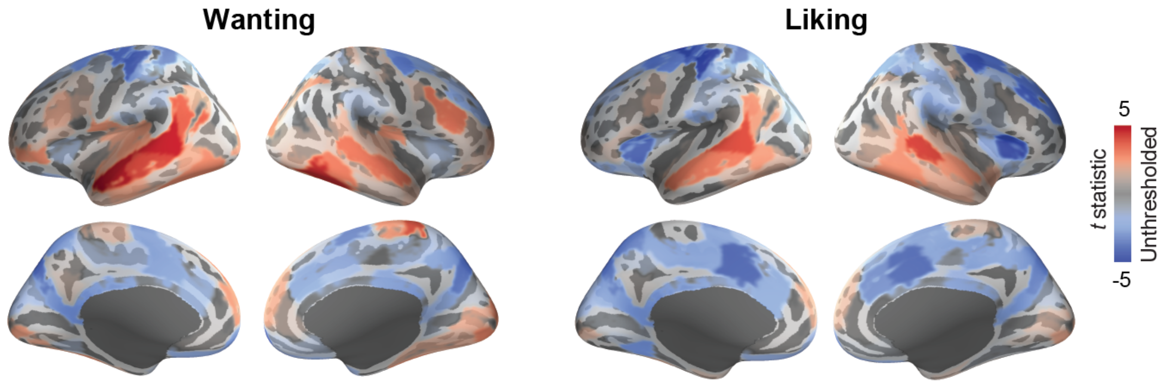


**Supplementary Fig. 2**. **Unthresholded cortical activation maps associated with wanting and liking during active information seeking, based on separate models.** Red and blue regions indicate positive and negative associations with the motivational dimension of interest, respectively. For each cortical parcel, we fit separate linear mixed-effects models relating event-wise mean neural activation to either wanting or liking. The resulting *t*-statistic maps are displayed on the inflated surface of the FreeSurfer template brain (fsaverage).

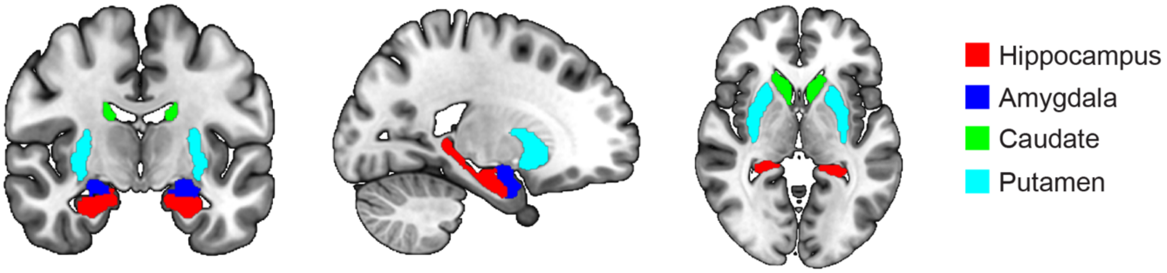


**Supplementary Fig. 3**. **Subcortical regions of interest (ROI).** Anatomically defined bilateral hippocampus, amygdala, caudate, and putamen ROIs are shown on coronal (left), sagittal (middle), and axial (right) slices of the MNI template brain.

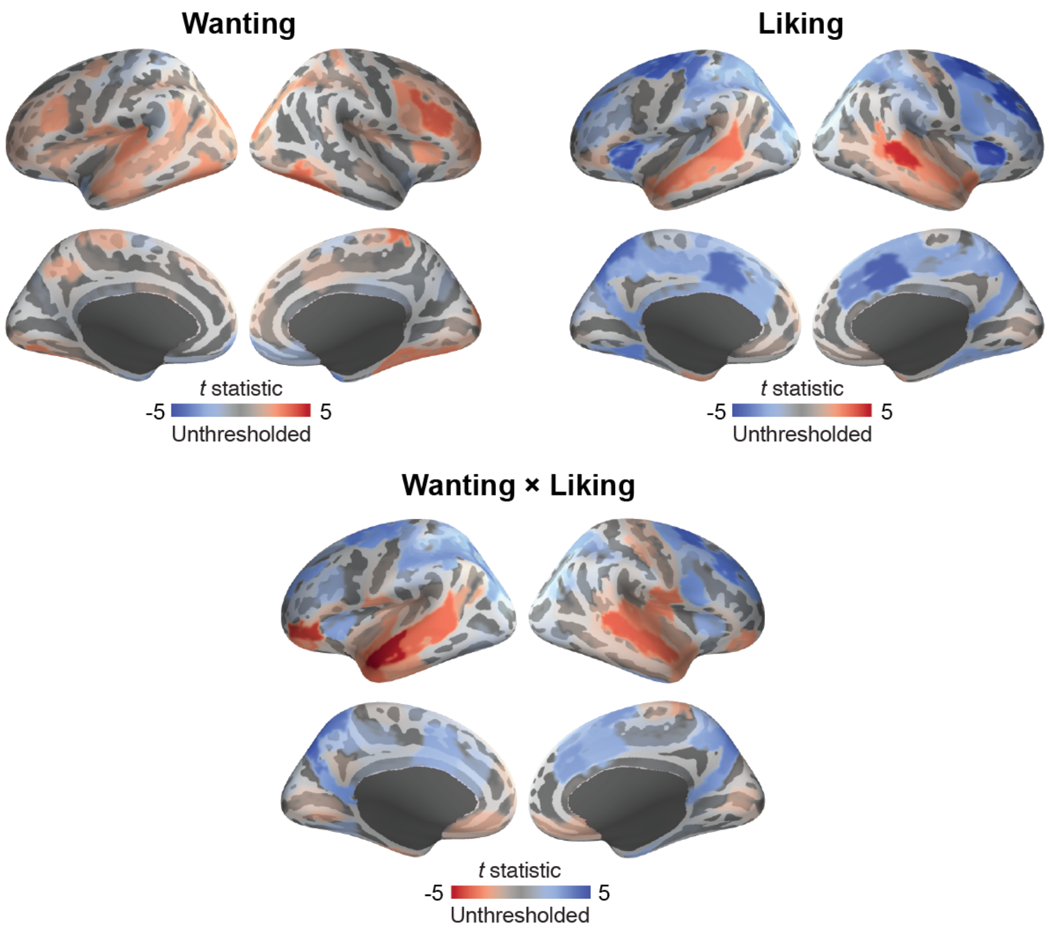


**Supplementary Fig. 4. Unthresholded cortical activation maps for wanting, liking, and their interaction during active information seeking.** Maps show *t*-statistics from a linear mixed-effects model relating event-wise mean neural activation to wanting, liking, and their interaction. For the wanting and liking maps, red and blue indicate positive and negative associations with the predictor of interest, respectively. For the interaction map, red indicates a relatively more positive effect of wanting when liking is lower, whereas blue indicates a relatively more positive effect of wanting when liking is higher. All maps are displayed on the inflated surface of the FreeSurfer template brain (fsaverage).

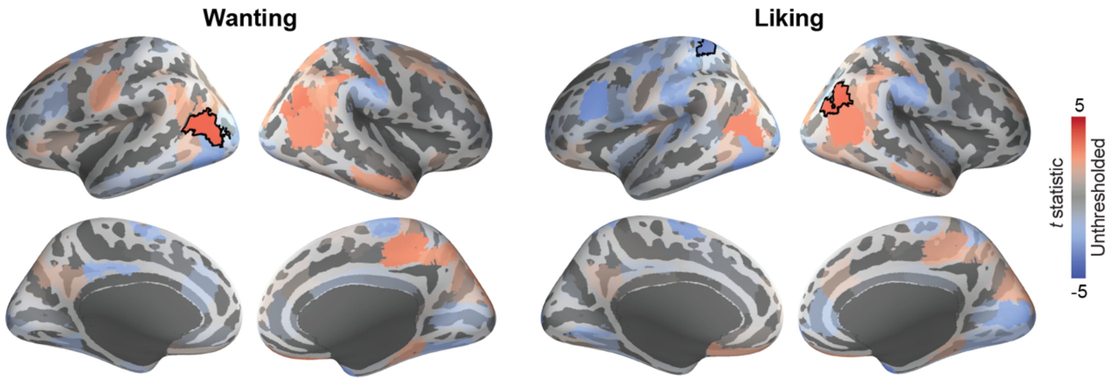


**Supplementary Fig. 5**. **Cortical regions in which event-wise seeker-viewer intersubject pattern correlation during encoding was associated with the pair's mean wanting (left panel) or liking (right panel) ratings of the events.** Red and blue regions indicate positive and negative associations, respectively, as estimated using linear mixed-effects models. The resulting unthresholded *t*-statistic maps are displayed on the inflated surfaces of the FreeSurfer template brain (fsaverage). Black outlines denote parcels that survived FDR correction across cortical parcels.

### SUPPLEMENTARY TABLES

**Supplementary Table 1. Group-level descriptive statistics for event-wise intrinsic motivation, stimulus properties, and web browsing actions (N = 54).**

| Variable* | Mean | SD | Range |
|---|---|---|---|
| Liking (1 = not at all, 5 = very much) | 3.18 | 0.68 | 1.79–4.87 |
| Wanting (1 = not at all, 5 = very much) | 3.61 | 0.65 | 1.83–5.00 |
| Audio amplitude (arbitrary unit) | 155.32 | 333.68 | 0.85–2223.15 |
| Brightness (0 = black, 255 = white) | 200.86 | 14.35 | 155.56–225.70 |
| Keypress (frequency) | 0.04 | 0.03 | 0.00–0.10 |
| Click (frequency) | 0.23 | 0.07 | 0.08–0.39 |
| Scroll (frequency) | 0.29 | 0.07 | 0.17–0.49 |
| Mouse movement (distance in pixel) | 218.92 | 87.81 | 59.64–467.69 |

*Values were first averaged across TRs within each event and then across events within each participant.

**Supplementary Table 2. One-sample *t*-tests of browsing action-motivation correlations against zero.**

| Correlated variables | Mean Fisher's *z* | SD | *t** | *df* | *p* (unc.) |
|---|---|---|---|---|---|
| Keypress-Liking | -0.06 | 0.12 | -3.60 | 52 | 0.001 |
| Keypress-Wanting | 0.03 | 0.12 | 1.78 | 51 | 0.082 |
| Click-Liking | -0.09 | 0.13 | -5.12 | 53 | <0.001 |
| Click-Wanting | -0.07 | 0.13 | -3.84 | 52 | <0.001 |
| Scroll-Liking | -0.02 | 0.15 | -1.23 | 53 | 0.225 |
| Scroll-Wanting | -0.01 | 0.12 | -0.70 | 52 | 0.486 |
| Mouse movement-Liking | -0.10 | 0.14 | -5.24 | 53 | <0.001 |
| Mouse movement- Wanting | -0.06 | 0.15 | -3.07 | 52 | 0.003 |

*Two-tailed tests

**Supplementary Table 3. Cortical parcels showing a significant main effect of wanting on neural activation during active web browsing ($q_{FDR} < 0.05$).**

| Hemisphere | Network | Parcel | $\beta$* | $t$ | $p$ (unc.) |
|---|---|---|---|---|---|
| Left | ContA | PFCd_1 | -0.009 | -2.93 | 0.003 |
| Left | ContC | Cingp_1 | -0.009 | -2.87 | 0.004 |
| Left | ContC | pCun_1 | -0.016 | -3.84 | <0.001 |
| Left | DefaultB | IPL_1 | 0.013 | 3.66 | <0.001 |
| Left | DefaultB | PFCd_2 | 0.011 | 3.12 | 0.002 |
| Left | DefaultB | PFCv_3 | 0.008 | 2.70 | 0.007 |
| Left | DefaultB | Temp_3 | 0.014 | 5.04 | <0.001 |
| Left | DefaultB | Temp_4 | 0.015 | 4.19 | <0.001 |
| Left | DefaultC | Rsp_1 | -0.009 | -2.77 | 0.006 |
| Left | DorsAttnA | SPL_2 | -0.015 | -3.60 | <0.001 |
| Left | DorsAttnA | TempOcc_1 | 0.007 | 2.71 | 0.007 |
| Left | DorsAttnA | TempOcc_2 | 0.011 | 2.78 | 0.006 |
| Left | DorsAttnB | PostC_3 | -0.012 | -3.46 | 0.001 |
| Left | SalVentAttnA | FrMed_2 | -0.008 | -2.67 | 0.008 |
| Left | SomMotA | SomMotA_2 | -0.010 | -2.66 | 0.008 |
| Left | SomMotA | SomMotA_3 | -0.019 | -4.34 | <0.001 |
| Left | SomMotA | SomMotA_4 | -0.013 | -3.81 | <0.001 |
| Left | SomMotA | SomMotA_5 | -0.011 | -3.15 | 0.002 |
| Left | SomMotA | SomMotA_6 | -0.013 | -3.86 | <0.001 |
| Left | TempPar | TempPar_1 | 0.017 | 4.76 | <0.001 |
| Left | TempPar | TempPar_2 | 0.013 | 3.73 | <0.001 |
| Right | ContA | PFCl_2 | 0.010 | 3.22 | 0.001 |
| Right | ContB | Temp_2 | 0.010 | 3.13 | 0.002 |
| Right | ContC | pCun_1 | -0.015 | -3.64 | <0.001 |
| Right | DorsAttnA | SPL_2 | -0.012 | -2.82 | 0.005 |
| Right | DorsAttnA | TempOcc_1 | 0.015 | 4.58 | <0.001 |
| Right | DorsAttnB | FEF_1 | -0.011 | -3.30 | 0.001 |
| Right | LimbicA | TempPole_4 | 0.007 | 2.71 | 0.007 |
| Right | LimbicB | OFC_4 | -0.012 | -4.19 | <0.001 |
| Right | SalVentAttnA | FrMed_2 | -0.010 | -3.35 | 0.001 |
| Right | SomMotA | SomMotA_8 | -0.008 | -2.77 | 0.006 |
| Right | SomMotA | SomMotA_9 | 0.010 | 3.93 | <0.001 |

| | | | | | |
|---|---|---|---|---|---|
| Right | TempPar | TempPar_1 | 0.009 | 3.31 | 0.001 |
| Right | TempPar | TempPar_2 | 0.011 | 3.22 | 0.001 |
| Right | VisCent | ExStr_5 | 0.011 | 2.77 | 0.006 |
| Right | VisCent | Striate_1 | 0.012 | 2.89 | 0.004 |

* Model: neural activation ~ wanting + run + screen brightness + audio amplitude + click + scroll + keypress + mouse movement + (1 | participant).

**Supplementary Table 4. Cortical parcels showing a significant main effect of liking on neural activation during active web browsing ($q_{FDR} < 0.05$).**

| Hemisphere | Network | Parcel | $\beta$* | $t$ | $p$ (unc.) |
|---|---|---|---|---|---|
| Left | ContA | Temp_1 | 0.008 | 2.58 | 0.010 |
| Left | ContC | Cingp_1 | -0.009 | -3.01 | 0.003 |
| Left | ContC | pCun_2 | -0.013 | -3.41 | <0.001 |
| Left | ContC | pCun_1 | -0.014 | -3.69 | <0.001 |
| Left | DefaultB | Temp_3 | 0.009 | 3.26 | 0.001 |
| Left | DefaultB | Temp_4 | 0.011 | 3.25 | 0.001 |
| Left | DefaultB | IPL_1 | 0.010 | 2.87 | 0.004 |
| Left | DefaultC | Rsp_1 | -0.008 | -2.72 | 0.007 |
| Left | DorsAttnA | SPL_2 | -0.016 | -4.03 | <0.001 |
| Left | DorsAttnB | FEF_1 | -0.009 | -3.38 | <0.001 |
| Left | DorsAttnB | PostC_3 | -0.013 | -3.73 | <0.001 |
| Left | SalVentAttnA | ParMed_1 | -0.007 | -2.62 | 0.009 |
| Left | SalVentAttnA | FrMed_2 | -0.008 | -2.73 | 0.006 |
| Left | SalVentAttnA | FrMed_1 | -0.012 | -4.35 | <0.001 |
| Left | SalVentAttnB | PFCmp_1 | -0.007 | -2.62 | 0.009 |
| Left | SalVentAttnB | Ins_1 | -0.011 | -4.29 | <0.001 |
| Left | SomMotA | SomMotA_1 | -0.007 | -2.52 | 0.012 |
| Left | SomMotA | SomMotA_2 | -0.010 | -2.76 | 0.006 |
| Left | SomMotA | SomMotA_3 | -0.022 | -5.25 | <0.001 |
| Left | SomMotA | SomMotA_4 | -0.014 | -4.29 | <0.001 |
| Left | SomMotA | SomMotA_5 | -0.009 | -2.60 | 0.009 |
| Left | SomMotA | SomMotA_6 | -0.016 | -5.00 | <0.001 |
| Left | TempPar | TempPar_1 | 0.015 | 4.29 | <0.001 |
| Left | VisPeri | ExStrInf_1 | -0.008 | -2.85 | 0.004 |
| Right | ContA | PFCd_1 | -0.012 | -3.91 | <0.001 |
| Right | ContB | Temp_2 | 0.008 | 2.78 | 0.006 |
| Right | ContB | PFCld_1 | -0.016 | -4.46 | <0.001 |
| Right | ContC | pCun_2 | -0.009 | -2.70 | 0.007 |
| Right | ContC | pCun_1 | -0.013 | -3.26 | 0.001 |
| Right | DefaultA | PFCd_1 | -0.011 | -3.85 | <0.001 |
| Right | DorsAttnA | TempOcc_1 | 0.008 | 2.68 | 0.007 |
| Right | DorsAttnA | SPL_4 | -0.010 | -2.66 | 0.008 |

| | | | | | |
|---|---|---|---|---|---|
| Right | DorsAttnA | SPL_2 | -0.015 | -3.54 | <0.001 |
| Right | DorsAttnB | FEF_1 | -0.015 | -4.71 | <0.001 |
| Right | LimbicB | OFC_4 | -0.008 | -2.94 | 0.003 |
| Right | SalVentAttnA | ParMed_2 | -0.008 | -2.73 | 0.006 |
| Right | SalVentAttnA | FrMed_2 | -0.008 | -2.81 | 0.005 |
| Right | SalVentAttnA | FrMed_1 | -0.010 | -4.21 | <0.001 |
| Right | SalVentAttnB | PFCmp_1 | -0.009 | -3.31 | <0.001 |
| Right | SalVentAttnB | PFCl_1 | -0.012 | -3.80 | <0.001 |
| Right | SalVentAttnB | Ins_2 | -0.016 | -5.16 | <0.001 |
| Right | SomMotA | SomMotA_3 | -0.007 | -2.65 | 0.008 |
| Right | SomMotA | SomMotA_5 | -0.010 | -2.80 | 0.005 |
| Right | SomMotA | SomMotA_8 | -0.010 | -3.55 | <0.001 |
| Right | TempPar | TempPar_1 | 0.007 | 2.68 | 0.007 |
| Right | TempPar | TempPar_2 | 0.015 | 4.45 | <0.001 |
| Right | TempPar | TempPar_3 | 0.010 | 2.73 | 0.007 |

* Model: neural activation ~ liking + run + screen brightness + audio amplitude + click + scroll + keypress + mouse movement + (1 | participant).

**Supplementary Table 5. Results of separate models examining associations between subcortical regional activation and wanting or liking during active web browsing.**

| Predictor | Region | $\beta$* | $t$ | $p$ (unc.) |
|---|---|---|---|---|
| Wanting | Hippocampus | 0.006 | 2.70 | 0.007 |
| Wanting | Amygdala | 0.005 | 2.00 | 0.045 |
| Wanting | Caudate | 0.006 | 2.18 | 0.029 |
| Wanting | Putamen | 0.002 | 0.64 | 0.524 |
| Liking | Hippocampus | 0.008 | 3.80 | <0.001 |
| Liking | Amygdala | 0.007 | 3.09 | 0.002 |
| Liking | Caudate | 0.002 | 0.88 | 0.379 |
| Liking | Putamen | -0.002 | -0.81 | 0.416 |

* Model: neural activation ~ motivation (wanting or liking) + run + screen brightness + audio amplitude + click + scroll + keypress + mouse movement + (1 | participant).

**Supplementary Table 6. Cortical parcels in which activation during active web browsing was significantly predicted by wanting, liking, or their interaction ($q_{FDR} < 0.05$).**

| Predictor | Hemisphere | Network | Parcel | *β** | *t* | *p* (unc.) |
|---|---|---|---|---|---|---|
| Liking | Left | ContA | PFCd_1 | -0.015 | -3.82 | <0.001 |
| Liking | Left | ContC | pCun_1 | -0.020 | -3.56 | <0.001 |
| Liking | Left | ContC | pCun_2 | -0.021 | -3.85 | <0.001 |
| Liking | Left | DefaultB | Temp_3 | 0.012 | 3.16 | 0.002 |
| Liking | Left | DefaultB | Temp_4 | 0.013 | 2.68 | 0.007 |
| Liking | Left | DefaultC | IPL_1 | -0.015 | -3.12 | 0.002 |
| Liking | Left | DorsAttnA | SPL_1 | -0.017 | -3.11 | 0.002 |
| Liking | Left | DorsAttnA | SPL_2 | -0.024 | -4.25 | <0.001 |
| Liking | Left | DorsAttnB | FEF_1 | -0.016 | -4.26 | <0.001 |
| Liking | Left | DorsAttnB | PostC_3 | -0.013 | -2.81 | 0.005 |
| Liking | Left | DorsAttnB | PostC_4 | -0.012 | -2.62 | 0.009 |
| Liking | Left | SalVentAttnA | FrMed_1 | -0.015 | -4.04 | <0.001 |
| Liking | Left | SalVentAttnA | ParMed_1 | -0.010 | -2.66 | 0.008 |
| Liking | Left | SalVentAttnB | Ins_1 | -0.018 | -4.78 | <0.001 |
| Liking | Left | SalVentAttnB | PFCl_1 | -0.011 | -2.74 | 0.006 |
| Liking | Left | SomMotA | SomMotA_3 | -0.022 | -3.83 | <0.001 |
| Liking | Left | SomMotA | SomMotA_4 | -0.017 | -3.61 | <0.001 |
| Liking | Left | SomMotA | SomMotA_6 | -0.020 | -4.50 | <0.001 |
| Liking | Left | TempPar | TempPar_1 | 0.016 | 3.38 | <0.001 |
| Liking | Left | VisCent | ExStr_5 | -0.019 | -3.50 | <0.001 |
| Liking | Left | VisPeri | ExStrInf_1 | -0.013 | -3.09 | 0.002 |
| Liking | Right | ContA | PFCd_1 | -0.021 | -4.72 | <0.001 |
| Liking | Right | ContB | PFCld_1 | -0.027 | -5.24 | <0.001 |
| Liking | Right | ContB | PFCld_2 | -0.013 | -2.90 | 0.004 |
| Liking | Right | ContC | pCun_1 | -0.016 | -2.78 | 0.005 |
| Liking | Right | ContC | pCun_2 | -0.013 | -2.64 | 0.008 |
| Liking | Right | DefaultA | PFCd_1 | -0.019 | -4.73 | <0.001 |
| Liking | Right | DorsAttnA | SPL_2 | -0.021 | -3.63 | <0.001 |
| Liking | Right | DorsAttnA | SPL_3 | -0.013 | -2.87 | 0.004 |
| Liking | Right | DorsAttnA | SPL_4 | -0.015 | -2.88 | 0.004 |
| Liking | Right | DorsAttnB | FEF_1 | -0.018 | -4.14 | <0.001 |
| Liking | Right | SalVentAttnA | FrMed_1 | -0.016 | -4.49 | <0.001 |
| Liking | Right | SalVentAttnA | Ins_1 | 0.008 | 2.82 | 0.005 |
| Liking | Right | SalVentAttnB | Ins_2 | -0.024 | -5.59 | <0.001 |
| Liking | Right | SalVentAttnB | PFCl_1 | -0.019 | -4.08 | <0.001 |
| Liking | Right | SalVentAttnB | PFCmp_1 | -0.014 | -3.52 | <0.001 |
| Liking | Right | SomMotA | SomMotA_8 | -0.013 | -3.22 | 0.001 |

| | | | | | | |
|---|---|---|---|---|---|---|
| Liking | Right | TempPar | TempPar_1 | 0.010 | 2.71 | 0.007 |
| Liking | Right | TempPar | TempPar_2 | 0.021 | 4.59 | <0.001 |
| Liking | Right | TempPar | TempPar_3 | 0.014 | 2.79 | 0.005 |
| Wanting | Right | ContA | PFCl_2 | 0.015 | 3.67 | <0.001 |
| Liking × Wanting | Left | ContA | IPS_1 | 0.009 | 2.86 | 0.004 |
| Liking × Wanting | Left | ContA | IPS_2 | 0.008 | 2.76 | 0.006 |
| Liking × Wanting | Left | ContA | IPS_3 | 0.009 | 3.04 | 0.002 |
| Liking × Wanting | Left | ContA | PFCd_1 | 0.008 | 3.38 | <0.001 |
| Liking × Wanting | Left | ContB | PFClv_1 | -0.012 | -4.35 | <0.001 |
| Liking × Wanting | Left | ContC | pCun_1 | 0.013 | 3.96 | <0.001 |
| Liking × Wanting | Left | ContC | pCun_2 | 0.011 | 3.52 | <0.001 |
| Liking × Wanting | Left | DefaultB | PFCv_2 | -0.006 | -3.05 | 0.002 |
| Liking × Wanting | Left | DefaultB | PFCv_3 | -0.011 | -4.33 | <0.001 |
| Liking × Wanting | Left | DefaultB | Temp_1 | -0.007 | -3.43 | <0.001 |
| Liking × Wanting | Left | DefaultB | Temp_3 | -0.012 | -5.45 | <0.001 |
| Liking × Wanting | Left | DefaultB | Temp_4 | -0.010 | -3.51 | <0.001 |
| Liking × Wanting | Left | DefaultC | IPL_1 | 0.008 | 2.81 | 0.005 |
| Liking × Wanting | Left | DorsAttnA | SPL_1 | 0.011 | 3.57 | <0.001 |
| Liking × Wanting | Left | DorsAttnA | SPL_2 | 0.014 | 4.33 | <0.001 |
| Liking × Wanting | Left | DorsAttnB | FEF_1 | 0.006 | 2.70 | 0.007 |
| Liking × Wanting | Left | SomMotA | SomMotA_4 | 0.007 | 2.70 | 0.007 |
| Liking × Wanting | Left | SomMotA | SomMotA_6 | 0.008 | 3.07 | 0.002 |
| Liking × Wanting | Left | TempPar | TempPar_1 | -0.010 | -3.43 | <0.001 |
| Liking × Wanting | Right | ContA | PFCd_1 | 0.010 | 3.99 | <0.001 |
| Liking × Wanting | Right | ContB | PFCld_1 | 0.011 | 3.75 | <0.001 |
| Liking × Wanting | Right | ContC | pCun_1 | 0.010 | 3.10 | 0.002 |
| Liking × Wanting | Right | DefaultA | PFCd_1 | 0.008 | 3.56 | <0.001 |
| Liking × Wanting | Right | DorsAttnA | SPL_2 | 0.011 | 3.22 | 0.001 |
| Liking × Wanting | Right | SalVentAttnA | FrMed_2 | 0.007 | 3.00 | 0.003 |
| Liking × Wanting | Right | SalVentAttnB | PFCmp_1 | 0.006 | 2.95 | 0.003 |
| Liking × Wanting | Right | SomMotB | Aud_2 | -0.008 | -3.03 | 0.002 |
| Liking × Wanting | Right | SomMotB | S2_1 | -0.006 | -2.87 | 0.004 |
| Liking × Wanting | Right | SomMotB | S2_3 | -0.008 | -2.98 | 0.003 |
| Liking × Wanting | Right | SomMotB | S2_4 | -0.007 | -3.02 | 0.003 |
| Liking × Wanting | Right | TempPar | TempPar_1 | -0.008 | -3.66 | <0.001 |
| Liking × Wanting | Right | TempPar | TempPar_2 | -0.010 | -3.71 | <0.001 |

* Model: neural activation ~ wanting + liking + wanting × liking + run + screen brightness + audio amplitude + click + scroll + keypress + mouse movement + (1 | participant).

**Supplementary Table 7. Results of models examining associations between subcortical regional activation and wanting, liking, and their interaction during active web browsing.**

| Predictor | Region | *β** | *t* | *p* (unc.) |
|---|---|---|---|---|
| Wanting | Hippocampus | -0.001 | -0.19 | 0.848 |
| Wanting | Amygdala | -0.001 | -0.40 | 0.688 |
| Wanting | Caudate | 0.007 | 2.06 | 0.039 |
| Wanting | Putamen | 0.005 | 1.35 | 0.177 |
| Liking | Hippocampus | 0.012 | 3.65 | <0.001 |
| Liking | Amygdala | 0.010 | 3.11 | 0.002 |
| Liking | Caudate | -0.002 | -0.60 | 0.549 |
| Liking | Putamen | -0.004 | -1.25 | 0.213 |
| Wanting × Liking | Hippocampus | -0.005 | -2.49 | 0.013 |
| Wanting × Liking | Amygdala | -0.004 | -2.11 | 0.035 |
| Wanting × Liking | Caudate | 0.001 | 0.25 | 0.805 |
| Wanting × Liking | Putamen | <0.001 | -0.09 | 0.932 |

* Model: neural activation ~ wanting + liking + wanting × liking + run + screen brightness + audio amplitude + click + scroll + keypress + mouse-movement amplitude + (1 | participant).

**Supplementary Table 8. Simple slope analyses of the interactive effects of wanting and liking on hippocampal and a right superior temporal sulcus parcel (TempPar_1) activation during active web browsing.**

| Region | Predictor | Moderator | Moderator level | $\beta$ | $t$ | $p$ (unc.) |
|---|---|---|---|---|---|---|
| Hippocampus | Liking | Wanting | Low (-1 SD) | 0.010 | 3.72 | <0.001 |
| Hippocampus | Liking | Wanting | High (+1 SD) | 0.004 | 1.08 | 0.280 |
| Hippocampus | Wanting | Liking | Low (-1 SD) | 0.004 | 1.36 | 0.175 |
| Hippocampus | Wanting | Liking | High (+1 SD) | -0.007 | -1.60 | 0.110 |
| Right TempPar_1 | Liking | Wanting | Low (-1 SD) | 0.014 | 3.24 | 0.001 |
| Right TempPar_1 | Liking | Wanting | High (+1 SD) | -0.004 | -1.09 | 0.274 |
| Right TempPar_1 | Wanting | Liking | Low (-1 SD) | 0.012 | 3.31 | <0.001 |
| Right TempPar_1 | Wanting | Liking | High (+1 SD) | -0.008 | -1.47 | 0.141 |

**Supplementary Table 9. Simple slope analyses of the interactive effects of wanting and liking on recall success across all events in the active information-seeking group.**

| Predictor | Moderator | Moderator level | $\beta$ | $z$ | $p$ (unc.) |
|---|---|---|---|---|---|
| Liking | Wanting | Low (-1 SD) | 0.30 | 7.699 | <0.001 |
| Liking | Wanting | High (+1 SD) | 0.10 | 2.998 | 0.003 |
| Wanting | Liking | Low (-1 SD) | 0.13 | 4.010 | <0.001 |
| Wanting | Liking | High (+1 SD) | -0.09 | -1.852 | 0.064 |

**Supplementary Table 10. Cortical parcels in which activation during active web browsing significantly predicted subsequent recall ($q_{FDR} < 0.05$).**

| Hemisphere | Network | Parcel | $\beta$* | z | *p* (unc.) |
|---|---|---|---|---|---|
| Left | ContA | IPS_1 | -0.18 | -2.63 | 0.008 |
| Left | ContA | IPS_2 | -0.25 | -3.48 | <0.001 |
| Left | ContA | IPS_3 | -0.20 | -2.81 | 0.005 |
| Left | ContA | PFCd_1 | -0.48 | -5.04 | <0.001 |
| Left | ContA | PFClv_1 | -0.41 | -4.49 | <0.001 |
| Left | ContB | PFClv_2 | -0.17 | -2.49 | 0.013 |
| Left | ContC | Cingp_1 | -0.32 | -3.97 | <0.001 |
| Left | ContC | pCun_1 | -0.38 | -5.80 | <0.001 |
| Left | ContC | pCun_2 | -0.39 | -5.64 | <0.001 |
| Left | DefaultB | PFCd_2 | 0.24 | 3.40 | <0.001 |
| Left | DefaultB | PFCv_2 | -0.45 | -4.31 | <0.001 |
| Left | DefaultB | Temp_2 | 0.27 | 2.65 | 0.008 |
| Left | DefaultB | Temp_3 | 0.45 | 4.67 | <0.001 |
| Left | DefaultB | Temp_4 | 0.33 | 4.35 | <0.001 |
| Left | DefaultC | IPL_1 | -0.25 | -3.34 | <0.001 |
| Left | DorsAttnA | SPL_2 | -0.36 | -5.27 | <0.001 |
| Left | DorsAttnB | FEF_1 | -0.39 | -3.84 | <0.001 |
| Left | DorsAttnB | PostC_1 | -0.36 | -4.76 | <0.001 |
| Left | DorsAttnB | PostC_2 | -0.28 | -3.94 | <0.001 |
| Left | DorsAttnB | PostC_3 | -0.27 | -3.46 | <0.001 |
| Left | DorsAttnB | PostC_4 | -0.23 | -2.78 | 0.005 |
| Left | LimbicA | TempPole_3 | -0.40 | -2.71 | 0.007 |
| Left | LimbicB | OFC_2 | -0.38 | -2.98 | 0.003 |
| Left | SalVentAttnA | FrMed_1 | -0.43 | -4.49 | <0.001 |
| Left | SalVentAttnA | FrMed_2 | -0.28 | -3.17 | 0.001 |
| Left | SalVentAttnA | FrOper_1 | -0.25 | -2.48 | 0.013 |
| Left | SalVentAttnA | ParMed_1 | -0.32 | -3.30 | <0.001 |
| Left | SalVentAttnA | ParOper_1 | -0.32 | -3.55 | <0.001 |
| Left | SalVentAttnB | Ins_1 | -0.42 | -4.29 | <0.001 |
| Left | SalVentAttnB | PFCl_1 | -0.32 | -3.60 | <0.001 |
| Left | SalVentAttnB | PFCmp_1 | -0.22 | -2.58 | 0.010 |
| Left | SomMotA | SomMotA_1 | -0.26 | -2.67 | 0.008 |

| | | | | | |
|---|---|---|---|---|---|
| Left | SomMotA | SomMotA_2 | -0.29 | -3.96 | <0.001 |
| Left | SomMotA | SomMotA_3 | -0.34 | -5.25 | <0.001 |
| Left | SomMotA | SomMotA_4 | -0.36 | -4.51 | <0.001 |
| Left | SomMotA | SomMotA_6 | -0.41 | -4.72 | <0.001 |
| Left | SomMotB | S2_3 | -0.22 | -2.75 | 0.006 |
| Left | TempPar | TempPar_1 | 0.35 | 4.70 | <0.001 |
| Left | TempPar | TempPar_2 | 0.31 | 4.04 | <0.001 |
| Left | VisCent | ExStr_3 | 0.18 | 2.85 | 0.004 |
| Left | VisCent | Striate_1 | 0.23 | 3.49 | <0.001 |
| Right | ContA | PFCd_1 | -0.35 | -4.16 | <0.001 |
| Right | ContA | PFCl_1 | -0.21 | -2.45 | 0.014 |
| Right | ContB | PFCld_1 | -0.22 | -3.14 | 0.002 |
| Right | ContB | Temp_2 | 0.23 | 2.78 | 0.005 |
| Right | ContC | Cingp_1 | -0.29 | -3.00 | 0.003 |
| Right | ContC | pCun_1 | -0.32 | -5.01 | <0.001 |
| Right | ContC | pCun_2 | -0.24 | -3.22 | 0.001 |
| Right | DefaultA | PFCd_1 | -0.33 | -3.61 | <0.001 |
| Right | DefaultC | Rsp_1 | -0.20 | -2.52 | 0.012 |
| Right | DorsAttnA | SPL_2 | -0.36 | -5.60 | <0.001 |
| Right | DorsAttnB | FEF_1 | -0.31 | -3.64 | <0.001 |
| Right | DorsAttnB | PostC_1 | -0.19 | -2.81 | 0.005 |
| Right | DorsAttnB | PostC_3 | -0.29 | -3.71 | <0.001 |
| Right | LimbicA | TempPole_1 | -0.34 | -2.49 | 0.013 |
| Right | LimbicB | OFC_1 | -0.42 | -3.98 | <0.001 |
| Right | LimbicB | OFC_4 | -0.24 | -2.52 | 0.012 |
| Right | SalVentAttnA | FrMed_1 | -0.42 | -3.95 | <0.001 |
| Right | SalVentAttnA | FrMed_2 | -0.38 | -3.96 | <0.001 |
| Right | SalVentAttnA | ParMed_2 | -0.34 | -3.79 | <0.001 |
| Right | SalVentAttnB | IPL_1 | -0.21 | -2.99 | 0.003 |
| Right | SalVentAttnB | Ins_2 | -0.25 | -2.93 | 0.003 |
| Right | SalVentAttnB | PFCl_1 | -0.36 | -4.59 | <0.001 |
| Right | SalVentAttnB | PFClv_1 | -0.22 | -2.86 | 0.004 |
| Right | SalVentAttnB | PFCmp_1 | -0.31 | -3.29 | 0.001 |
| Right | SomMotA | SomMotA_3 | -0.25 | -2.49 | 0.013 |
| Right | SomMotA | SomMotA_8 | -0.36 | -3.89 | <0.001 |
| Right | SomMotB | Aud_1 | 0.21 | 2.47 | 0.014 |

| | | | | | |
|---|---|---|---|---|---|
| Right | TempPar | TempPar_1 | 0.27 | 2.68 | 0.007 |
| Right | TempPar | TempPar_2 | 0.21 | 2.68 | 0.007 |
| Right | VisCent | Striate_1 | 0.24 | 3.50 | <0.001 |

* Model: recall success ~ activation + run + screen brightness + audio amplitude + click + scroll + keypress + mouse movement + (1 | participant).

**Supplementary Table 11. Results of models examining associations between subcortical regional activation during active web browsing and subsequent recall.**

| Region | $\beta$* | z | $p$ (unc.) |
|---|---|---|---|
| Amygdala | 0.259 | 2.320 | 0.020 |
| Caudate | 0.018 | 0.191 | 0.848 |
| Hippocampus | 0.186 | 1.686 | 0.092 |
| Putamen | -0.213 | -2.162 | 0.031 |

* Model: recall success ~ activation + run + screen brightness + audio amplitude + click + scroll + keypress + mouse movement + (1 | participant).

**Supplementary Table 12. Cortical parcels showing greater encoding-retrieval pattern similarity for matched than mismatched event pairs in the active information-seeking group ($q_{FDR} < 0.05$).**

| Hemisphere | Network | Parcel | *t** | *p (*unc.*)* |
|---|---|---|---|---|
| Left | ContA | IPS_1 | 3.29 | 0.002 |
| Left | ContA | PFCl_1 | 4.98 | <0.001 |
| Left | ContA | PFCl_2 | 4.43 | <0.001 |
| Left | ContA | PFCl_3 | 3.36 | 0.002 |
| Left | ContA | PFClv_1 | 2.89 | 0.006 |
| Left | ContA | Temp_1 | 5.10 | <0.001 |
| Left | ContB | IPL_1 | 4.47 | <0.001 |
| Left | ContB | PFCl_1 | 3.25 | 0.002 |
| Left | ContB | Temp_1 | 3.22 | 0.002 |
| Left | ContC | pCun_1 | 3.07 | 0.004 |
| Left | DefaultA | IPL_1 | 5.83 | <0.001 |
| Left | DefaultA | pCunPCC_1 | 4.79 | <0.001 |
| Left | DefaultA | pCunPCC_2 | 3.94 | <0.001 |
| Left | DefaultA | pCunPCC_3 | 4.07 | <0.001 |
| Left | DefaultA | PFCd_1 | 5.27 | <0.001 |
| Left | DefaultB | IPL_1 | 3.00 | 0.004 |
| Left | DefaultB | PFCd_1 | 2.97 | 0.005 |
| Left | DefaultB | PFCd_2 | 4.70 | <0.001 |
| Left | DefaultB | PFCd_4 | 5.05 | <0.001 |
| Left | DefaultB | PFCv_4 | 4.14 | <0.001 |
| Left | DefaultB | Temp_1 | 3.35 | 0.002 |
| Left | DefaultB | Temp_2 | 3.29 | 0.002 |
| Left | DefaultB | Temp_3 | 5.17 | <0.001 |
| Left | DefaultB | Temp_4 | 5.46 | <0.001 |
| Left | DefaultC | IPL_1 | 5.80 | <0.001 |
| Left | DefaultC | PHC_1 | 3.80 | <0.001 |
| Left | DefaultC | Rsp_1 | 5.40 | <0.001 |
| Left | DorsAttnA | TempOcc_1 | 3.98 | <0.001 |
| Left | DorsAttnA | TempOcc_2 | 3.49 | 0.001 |
| Left | SalVentAttnA | FrOper_1 | 2.69 | 0.010 |
| Left | SalVentAttnA | FrOper_2 | 3.64 | 0.001 |
| Left | SalVentAttnA | ParOper_1 | 2.59 | 0.013 |

| | | | | |
|---|---|---|---|---|
| Left | SalVentAttnB | IPL_1 | 2.58 | 0.013 |
| Left | SalVentAttnB | PFCl_1 | 3.01 | 0.004 |
| Left | SalVentAttnB | PFCmp_1 | 2.52 | 0.015 |
| Left | SomMotB | Cent_1 | 2.49 | 0.016 |
| Left | TempPar | TempPar_1 | 4.44 | <0.001 |
| Left | TempPar | TempPar_2 | 3.61 | 0.001 |
| Left | VisPeri | ExStrInf_3 | 3.00 | 0.004 |
| Right | ContA | Cingm_1 | 4.08 | <0.001 |
| Right | ContA | PFCd_1 | 3.86 | <0.001 |
| Right | ContA | PFCl_2 | 3.42 | 0.001 |
| Right | ContB | IPL_1 | 4.06 | <0.001 |
| Right | ContB | PFCld_1 | 2.66 | 0.011 |
| Right | ContB | PFCld_2 | 4.85 | <0.001 |
| Right | ContB | PFCld_3 | 3.67 | 0.001 |
| Right | ContB | PFClv_1 | 3.05 | 0.004 |
| Right | ContB | PFCmp_1 | 4.10 | <0.001 |
| Right | ContB | Temp_1 | 3.24 | 0.002 |
| Right | ContC | pCun_2 | 3.55 | 0.001 |
| Right | DefaultA | IPL_1 | 4.24 | <0.001 |
| Right | DefaultA | pCunPCC_1 | 3.73 | 0.001 |
| Right | DefaultA | PFCd_1 | 3.97 | <0.001 |
| Right | DefaultB | PFCd_1 | 3.32 | 0.002 |
| Right | DefaultB | PFCv_1 | 3.81 | <0.001 |
| Right | DefaultC | IPL_1 | 5.16 | <0.001 |
| Right | DefaultC | Rsp_1 | 3.47 | 0.001 |
| Right | DorsAttnA | ParOcc_1 | 3.09 | 0.003 |
| Right | SalVentAttnA | FrMed_2 | 2.57 | 0.014 |
| Right | SalVentAttnB | Ins_2 | 2.98 | 0.005 |
| Right | SalVentAttnB | PFCl_1 | 2.60 | 0.013 |
| Right | SalVentAttnB | PFClv_1 | 3.51 | 0.001 |
| Right | SalVentAttnB | PFCmp_1 | 3.50 | 0.001 |
| Right | TempPar | TempPar_1 | 2.87 | 0.006 |
| Right | TempPar | TempPar_2 | 3.07 | 0.004 |
| Right | TempPar | TempPar_3 | 3.14 | 0.003 |

* Two-tailed paired *t*-tests, degree of freedom = 46

**Supplementary Table 13. Comparisons of survey measures between the active information-seeking group (N = 54) and the passive viewing group (N = 23).**

| Variable | Seeker mean (SD) | Viewer mean (SD) | Test statistic* | *p* (unc.) | Effect size* | $q_{FDR}$ |
|---|---|---|---|---|---|---|
| Big 5 - Openness | 36.19 (7.29) | 39.61 (5.57) | -2.01 | 0.048 | -0.50 | 0.371 |
| Big 5 - Conscientiousness | 32.50 (6.58) | 33.83 (5.47) | -0.85 | 0.398 | -0.21 | 0.985 |
| Big 5 - Extraversion | 26.22 (6.75) | 26.48 (5.78) | -0.16 | 0.874 | -0.04 | 0.985 |
| Big 5 - Agreeableness | 34.06 (5.62) | 33.96 (4.94) | 0.07 | 0.942 | 0.02 | 0.985 |
| Big 5 - Neuroticism | 23.22 (6.32) | 22.65 (7.53) | 0.34 | 0.733 | 0.08 | 0.985 |
| Mind wandering | 18.89 (5.06) | 18.57 (5.22) | 0.25 | 0.800 | 0.06 | 0.985 |
| Internet addiction | 44.02 (13.83) | 41.65 (13.26) | 0.70 | 0.489 | 0.17 | 0.985 |
| ADHD | 8.76 (4.23) | 8.22 (4.33) | 0.51 | 0.611 | 0.13 | 0.985 |
| Curiosity | 34.11 (6.93) | 37.74 (8.43) | -1.97 | 0.053 | -0.49 | 0.371 |
| DASS - Stress | 12.41 (9.85) | 11.39 (9.29) | 0.42 | 0.675 | 0.10 | 0.985 |
| DASS - Anxiety | 8.37 (7.49) | 7.30 (8.06) | 0.56 | 0.578 | 0.14 | 0.985 |
| DASS - Depression | 9.41 (9.91) | 9.30 (11.00) | 0.04 | 0.968 | 0.01 | 0.985 |
| Age | 23.30 (4.70) | 21.2 (4.90) | 1.72 | 0.090 | 0.42 | 0.420 |
| Gender | Seeker N (%) | Viewer N (%) | 0.03 | 0.985 | 0.02 | 0.985 |
| Female | 30 (55.6%) | 13 (56.5%) | | | | |
| Male | 22 (40.7%) | 9 (39.1%) | | | | |
| Nonbinary | 2 (3.7%) | 1 (4.3%) | | | | |

* Test statistics were a $\chi^2$ statistic ($df = 2$) for the gender variable and $t$ statistics ($df = 75$) for all other variables. Effect sizes were Cramérs $V$ for the gender variable and Cohen's $d$ for all other variables.

**Supplementary Table 14. Comparisons of survey measures between viewers of Video 1 (N = 11) and viewers of Video 2 (N = 12).**

| Variable | Video 1 mean (SD) | Video 2 mean (SD) | Test statistic* | *p* (unc.) | Effect size* | $q_{FDR}$ |
|---|---|---|---|---|---|---|
| Big 5 - Openness | 40.18 (1.54) | 39.08 (1.77) | 0.47 | 0.645 | 0.19 | 0.754 |
| Big 5 - Conscientiousness | 36.45 (1.42) | 31.42 (1.48) | 2.45 | 0.023 | 1.02 | 0.222 |
| Big 5 - Extraversion | 26.55 (1.59) | 26.42 (1.86) | 0.05 | 0.959 | 0.02 | 0.959 |
| Big 5 - Agreeableness | 34.09 (1.09) | 33.83 (1.75) | 0.12 | 0.902 | 0.05 | 0.959 |
| Big 5 - Neuroticism | 21.82 (2.52) | 23.42 (2.01) | -0.5 | 0.626 | -0.21 | 0.754 |
| Mind wandering | 16.55 (1.59) | 20.42 (1.34) | -1.86 | 0.077 | -0.78 | 0.222 |
| Internet addiction | 37.36 (4.18) | 45.58 (3.44) | -1.52 | 0.145 | -0.64 | 0.337 |
| ADHD | 6.55 (1.38) | 9.75 (1.05) | -1.85 | 0.079 | -0.78 | 0.222 |
| Curiosity | 41.09 (2.56) | 34.67 (2.14) | 1.92 | 0.069 | 0.81 | 0.222 |
| DASS - Stress | 9.09 (2.69) | 13.50 (2.74) | -1.15 | 0.264 | -0.48 | 0.528 |
| DASS - Anxiety | 6.00 (2.21) | 8.50 (2.55) | -0.74 | 0.467 | -0.31 | 0.726 |
| DASS - Depression | 4.91 (2.25) | 13.33 (3.59) | -1.99 | 0.062 | -0.81 | 0.222 |
| Age | 20.73 (0.73) | 21.67 (1.87) | -0.47 | 0.646 | -0.19 | 0.754 |
| Gender | Video 1 N (%) | Video 2 N (%) | 2.04 | 0.361 | 0.30 | 0.632 |
| Female | 7 (63.6%) | 6 (50%) | | | | |
| Male | 3 (27.3%) | 6 (50%) | | | | |
| Nonbinary | 1 (9%) | 0 (0%) | | | | |

* Test statistics were a $\chi^2$ statistic (*df* = 2) for the gender variable and *t* statistics (*df* = 21) for all other variables. Effect sizes were Cramérs *V* for the gender variable and Cohen's *d* for all other variables.

**Supplementary Table 15. Behavioral comparisons between the active information-seeking group and the passive viewing groups for Video 1 and Video 2.**

| Variable | Video | Seeker mean (SD) | Viewer mean (SD) | *df* | *t** | *p* (unc.) |
|---|---|---|---|---|---|---|
| Wanting | Video 1 | 3.61 (0.65) | 2.80 (0.54) | 16.48 | 4.37 | <0.001 |
| Wanting | Video 2 | 3.61 (0.65) | 2.36 (0.46) | 21.88 | 7.83 | <0.001 |
| Liking | Video 1 | 3.18 (0.68) | 2.84 (0.62) | 15.39 | 1.61 | 0.128 |
| Liking | Video 2 | 3.18 (0.68) | 2.28 (0.39) | 28.16 | 6.13 | <0.001 |
| Wanting-Liking correlation† | Video 1 | 0.67 (0.56) | 1.44 (0.56) | 14.52 | -4.16 | 0.001 |
| Wanting-Liking correlation† | Video 2 | 0.67 (0.56) | 1.31 (0.66) | 14.81 | -3.09 | 0.008 |
| Memory performance | Video 1 | 0.59 (0.16) | 0.40 (0.21) | 12.47 | 2.86 | 0.014 |
| Memory performance | Video 2 | 0.59 (0.15) | 0.64 (0.16) | 16.4 | -0.98 | 0.340 |

* Group differences were evaluated using Welch's independent-samples *t*-tests to account for unequal variances and sample sizes.

† Fisher's *z*-transformed Pearson correlation across events.

**Supplementary Table 16. Behavioral comparisons between viewers and the corresponding seeker who created the video they watched.**

| Variable | Video | Seeker | Viewer mean (SD) | *df* | *t** | *p* (unc.) |
|---|---|---|---|---|---|---|
| Wanting | Video 1 | 3.46 | 2.80 (0.54) | 10 | 4.01 | 0.002 |
| Wanting | Video 2 | 3.77 | 2.36 (0.46) | 11 | 10.57 | <0.001 |
| Liking | Video 1 | 3.04 | 2.84 (0.62) | 10 | 1.06 | 0.314 |
| Liking | Video 2 | 3.74 | 2.28 (0.39) | 11 | 12.91 | <0.001 |
| Wanting-Liking correlation† | Video 1 | 0.56 | 1.44 (0.56) | 10 | -5.21 | <0.001 |
| Wanting-Liking correlation† | Video 2 | 1.59 | 1.31 (0.66) | 11 | 1.50 | 0.161 |
| Memory performance | Video 1 | 0.52 | 0.40 (0.21) | 10 | 1.82 | 0.098 |
| Memory performance | Video 2 | 0.59 | 0.64 (0.16) | 11 | -1.09 | 0.300 |

* Two-tailed one-sample *t*-tests.
† Fisher's *z*-transformed Pearson correlation across events.

**Supplementary Table 17. Cortical parcels showing a significant effect of pair-level wanting on viewer-viewer intersubject neural pattern similarity during encoding ($q_{FDR} < 0.05$).**

| Hemisphere | Network | Parcel | *β** | *t* | *p* (unc.) |
|---|---|---|---|---|---|
| Left | ContA | Temp_1 | 0.005 | 2.97 | 0.003 |
| Left | ContC | pCun_1 | 0.005 | 3.04 | 0.002 |
| Left | DefaultA | IPL_1 | 0.005 | 3.29 | <0.001 |
| Left | DefaultA | pCunPCC_1 | 0.004 | 3.46 | <0.001 |
| Left | DefaultB | PFCd_3 | 0.007 | 5.14 | <0.001 |
| Left | DefaultB | PFCd_4 | 0.005 | 3.51 | <0.001 |
| Left | DefaultB | PFCv_3 | 0.007 | 4.09 | <0.001 |
| Left | DefaultB | Temp_1 | 0.004 | 4.36 | <0.001 |
| Left | DefaultB | Temp_3 | 0.003 | 2.70 | 0.007 |
| Left | DefaultC | IPL_1 | 0.006 | 3.57 | <0.001 |
| Left | DefaultC | PHC_1 | 0.004 | 3.89 | <0.001 |
| Left | DorsAttnA | TempOcc_2 | -0.004 | -2.98 | 0.003 |
| Left | SalVentAttnA | FrMed_2 | -0.004 | -2.60 | 0.009 |
| Left | SalVentAttnA | Ins_1 | 0.003 | 3.28 | 0.001 |
| Left | SalVentAttnB | PFCl_1 | 0.003 | 2.90 | 0.004 |
| Left | SomMotB | Aud_2 | -0.003 | -2.95 | 0.003 |
| Left | SomMotB | S2_1 | 0.004 | 3.31 | <0.001 |
| Left | VisCent | ExStr_2 | -0.007 | -4.64 | <0.001 |
| Left | VisCent | Striate_1 | -0.004 | -2.57 | 0.010 |
| Left | VisPeri | ExStrInf_1 | 0.006 | 4.92 | <0.001 |
| Left | VisPeri | StriCal_1 | -0.005 | -4.33 | <0.001 |
| Right | ContA | IPS_1 | 0.007 | 4.61 | <0.001 |
| Right | ContA | IPS_2 | 0.005 | 3.05 | 0.002 |
| Right | ContA | PFCd_1 | 0.006 | 4.42 | <0.001 |
| Right | ContB | IPL_1 | 0.005 | 2.99 | 0.003 |
| Right | ContB | IPL_2 | 0.005 | 2.87 | 0.004 |
| Right | ContB | PFCld_2 | 0.005 | 3.70 | <0.001 |
| Right | ContB | PFCld_3 | 0.006 | 4.14 | <0.001 |
| Right | ContB | Temp_1 | 0.005 | 4.34 | <0.001 |
| Right | ContC | Cingp_1 | 0.007 | 4.84 | <0.001 |
| Right | ContC | pCun_1 | 0.007 | 4.41 | <0.001 |
| Right | ContC | pCun_2 | 0.005 | 3.41 | <0.001 |
| Right | DefaultA | IPL_1 | 0.006 | 4.17 | <0.001 |
| Right | DefaultA | PFCd_1 | 0.004 | 3.10 | 0.002 |
| Right | DefaultA | pCunPCC_1 | 0.004 | 3.56 | <0.001 |
| Right | DefaultB | AntTemp_1 | 0.004 | 3.77 | <0.001 |

| | | | | | |
|---|---|---|---|---|---|
| Right | DefaultB | PFCv_1 | 0.003 | 2.59 | 0.010 |
| Right | DefaultC | IPL_1 | 0.013 | 7.18 | <0.001 |
| Right | DefaultC | PHC_1 | 0.005 | 4.77 | <0.001 |
| Right | DorsAttnA | ParOcc_1 | 0.005 | 3.84 | <0.001 |
| Right | DorsAttnA | SPL_1 | 0.006 | 4.35 | <0.001 |
| Right | DorsAttnA | SPL_4 | 0.003 | 2.54 | 0.011 |
| Right | LimbicA | TempPole_1 | 0.003 | 2.78 | 0.005 |
| Right | LimbicA | TempPole_2 | 0.004 | 3.75 | <0.001 |
| Right | TempPar | TempPar_1 | 0.004 | 3.51 | <0.001 |
| Right | VisCent | ExStr_2 | -0.006 | -3.66 | <0.001 |
| Right | VisCent | ExStr_4 | -0.004 | -3.00 | 0.003 |
| Right | VisPeri | ExStrSup_3 | 0.005 | 2.93 | 0.003 |
| Right | VisPeri | StriCal_1 | -0.006 | -4.69 | <0.001 |

* Model: intersubject pattern similarity ~ pair-level mean wanting + video + run + audio amplitude + screen brightness + (1 | pair ID), where the pair ID indexed the unordered viewer-viewer pair.

**Supplementary Table 18. Cortical parcels showing a significant effect of pair-level liking on viewer-viewer intersubject neural pattern similarity during encoding ($q_{FDR} < 0.05$).**

| Hemisphere | Network | Parcel | *β** | *t* | *p* (unc.) |
|---|---|---|---|---|---|
| Left | VisCent | ExStr_2 | -0.008 | -4.85 | <0.001 |
| Left | VisPeri | ExStrInf_1 | 0.006 | 4.44 | <0.001 |
| Left | VisPeri | StriCal_1 | -0.006 | -4.70 | <0.001 |
| Left | SomMotB | Aud_2 | -0.004 | -3.48 | <0.001 |
| Left | SomMotB | S2_1 | 0.005 | 4.17 | <0.001 |
| Left | SomMotB | S2_3 | -0.004 | -2.77 | 0.006 |
| Left | DorsAttnA | TempOcc_1 | -0.003 | -2.66 | 0.008 |
| Left | SalVentAttnA | Ins_1 | 0.003 | 3.29 | 0.001 |
| Left | SalVentAttnA | FrMed_2 | -0.004 | -2.57 | 0.010 |
| Left | SalVentAttnB | PFCl_1 | 0.003 | 2.87 | 0.004 |
| Left | ContA | Temp_1 | 0.005 | 2.71 | 0.007 |
| Left | ContC | pCun_1 | 0.005 | 3.23 | 0.001 |
| Left | DefaultA | IPL_1 | 0.004 | 2.62 | 0.009 |
| Left | DefaultA | pCunPCC_1 | 0.004 | 3.27 | 0.001 |
| Left | DefaultB | Temp_1 | 0.003 | 3.40 | <0.001 |
| Left | DefaultB | PFCd_2 | 0.003 | 2.60 | 0.009 |
| Left | DefaultB | PFCd_3 | 0.006 | 3.92 | <0.001 |
| Left | DefaultB | PFCd_4 | 0.005 | 3.44 | <0.001 |
| Left | DefaultB | PFCv_2 | 0.005 | 3.05 | 0.002 |
| Left | DefaultB | PFCv_3 | 0.005 | 3.15 | 0.002 |
| Left | DefaultC | IPL_1 | 0.005 | 3.04 | 0.002 |
| Left | DefaultC | Rsp_1 | 0.004 | 2.79 | 0.005 |
| Left | DefaultC | PHC_1 | 0.006 | 4.90 | <0.001 |
| Right | VisCent | Striate_1 | -0.004 | -2.67 | 0.008 |
| Right | VisPeri | StriCal_1 | -0.007 | -5.07 | <0.001 |
| Right | DorsAttnA | ParOcc_1 | 0.007 | 5.02 | <0.001 |
| Right | DorsAttnA | SPL_1 | 0.006 | 3.90 | <0.001 |
| Right | SalVentAttnB | IPL_1 | 0.005 | 2.80 | 0.005 |
| Right | SalVentAttnB | PFCl_1 | 0.004 | 2.99 | 0.003 |
| Right | LimbicB | OFC_1 | 0.003 | 3.11 | 0.002 |
| Right | LimbicA | TempPole_2 | 0.005 | 4.19 | <0.001 |
| Right | ContA | IPS_1 | 0.005 | 2.83 | 0.005 |
| Right | ContA | PFCd_1 | 0.004 | 3.16 | 0.002 |
| Right | ContB | Temp_1 | 0.004 | 3.26 | 0.001 |
| Right | ContB | IPL_1 | 0.004 | 2.64 | 0.008 |
| Right | ContB | IPL_2 | 0.007 | 3.71 | <0.001 |

| | | | | | |
|---|---|---|---|---|---|
| Right | ContB | PFCld_2 | 0.004 | 2.91 | 0.004 |
| Right | ContB | PFCld_3 | 0.006 | 3.99 | <0.001 |
| Right | ContC | pCun_1 | 0.006 | 3.78 | <0.001 |
| Right | ContC | Cingp_1 | 0.006 | 4.41 | <0.001 |
| Right | DefaultA | IPL_1 | 0.007 | 4.74 | <0.001 |
| Right | DefaultA | PFCd_1 | 0.004 | 2.94 | 0.003 |
| Right | DefaultA | pCunPCC_1 | 0.005 | 3.72 | <0.001 |
| Right | DefaultB | AntTemp_1 | 0.003 | 2.57 | 0.010 |
| Right | DefaultC | IPL_1 | 0.014 | 7.51 | <0.001 |
| Right | DefaultC | PHC_1 | 0.006 | 5.26 | <0.001 |

*Model: intersubject pattern similarity ~ pair-level mean liking + video + run + audio amplitude + screen brightness + (1 | pair ID), where the pair ID indexed the unordered viewer-viewer pair.
